# A primordial TFEB-TGFβ signaling axis systemically regulates diapause and stem cell longevity

**DOI:** 10.1101/2023.10.06.561181

**Authors:** Tim J. Nonninger, Jennifer Mak, Birgit Gerisch, Valentina Ramponi, Kazuto Kawamura, Klara Schilling, Christian Latza, Jonathan Kölschbach, Roberto Ripa, Manuel Serrano, Adam Antebi

**Author notes:** Co-equal contribution (Tim J. Nonninger, Jennifer Mak, Birgit Gerisch).

## Abstract

Fasting/refeeding enhances animal health and lifespan across taxa. *C. elegans* can endure months of fasting in adult reproductive diapause (ARD) and upon refeeding, regenerate and reproduce. *hlh-30/TFEB* is an ARD master regulator whose mutants live mere days in ARD and don’t recover with refeeding. Here we find that downregulation of TGFβ signaling bypasses *hlh-30* collapse, and restores recovery, germline stem cell proliferation and reproductive competence. Upon fasting, HLH-30/TFEB(+) downregulates TGFβ in sensory neurons, to inhibit Notch and promote reproductive quiescence in the germline. Upon refeeding, these pathways are upregulated to activate stem cells and promote reproduction. *hlh-30* loss induces a senescent-like DNA damage, immune and growth metabolic signature reversed by inhibiting TGFβ signaling. TFEB’s role is conserved in mammalian diapause models, including mouse embryonic and human cancer diapause. Thus, TFEB-TGFβ axis relays systemic signals matching nutrient supply with growth signaling, to regulate stem cell longevity, senescence and regeneration across species.

## INTRODUCTION

During food scarcity or environmental stress, organisms throughout the tree of life can persist in long-lived states of quiescence such as diapause and torpor that enable them to outlast adversity until conditions improve.^1,2^ Even animals without explicit states of diapause, such as humans, globally remodel metabolism in response to fasting/refeeding and can enter hypometabolism. We can thus be thought of as cellular mosaics comprised of dormant diapause-like cells, and actively growing or dividing non-diapause cells,^3,4^ and this balance ensures stem cell longevity, tissue homeostasis and organismal lifespan^3,5,6^. In pathological cases, such mechanisms can be co-opted by tumor cells that enter diapause-like states to escape immune surveillance and resist chemotherapy, only to emerge later and develop into full blown cancer.^7,8^

A common feature of quiescence is the induction of resilience mechanisms that enable cellular and organismal survival.^9,10^ Notably, such mechanisms also impact overall lifespan, and many of the same molecular processes regulate organismal longevity across taxa.^1^ A return to favorable environments and nutrient-replete conditions triggers exit from quiescence. In the process, organisms often repair cumulative damage, and reconstitute somatic tissues and stem cell pools in a remarkable process of regeneration.^11^ Fasting/refeeding and quiescence/activation rely on metabolic flexibility to induce cellular plasticity and promote longevity.^3,5,6^ However, the molecular mechanisms underlying this remarkable plasticity and its decline with aging remain elusive.

Diapause is an extreme form of prolonged fasting and can serve as a model of quiescence from cellular to organismal levels. Molecular genetic dissection of the *C. elegans* dauer diapause^12^, for example, led to the seminal discovery of conserved nutrient sensing and growth signaling pathways such as insulin/IGF signaling and mTOR as conserved regulators of metazoan lifespan.^13,14^ Aside from dauer diapause, *C. elegans* can enter several other quiescent states during its life cycle, including the *C. elegans* adult reproductive diapause (ARD), a largely unstudied state of quiescence triggered in response to late larval stage starvation.^5,15^ Such animals exhibit adult features and live over 2 months without food, nearly 3 times longer than *ad libitum (AL)* fed adults.^16^ Upon refeeding, animals reactivate germline stem cells (GSC), regenerate tissues including soma and germline, reproduce and live normal lifespans, revealing extraordinary survivorship and rejuvenation in the adult animal. Our laboratory recently identified HLH-30/TFEB transcription factor as a master regulator of ARD, whose mutation drastically reduces survivorship to 5-10 days, and results in a failure to recover and produce progeny, reflecting GSC dysfunction.^16^ *hlh-30* mutants also exhibit a rapid shrinkage of body length, loss of body fat, and fragmented mitochondria within ARD. HLH-30/TFEB’s role is especially important to adult diapause, since *hlh-30* mutation disrupts ARD survival, metabolism and recovery, but has little effect on dauer diapause.

TFEB is best known as a regulator of autophagy and lysosome biogenesis^17–19^, yet mutations affecting these processes have surprisingly little impact on ARD survival.^16^ This raises the question: what mechanisms act downstream of HLH-30/TFEB to promote diapause survivorship and GSC longevity? Hence, we carried out unbiased suppressor screens, and unexpectedly found mutations in multiple components of transforming growth factor β (TGFβ) signaling that bypass *hlh-30* collapse, and stimulate rejuvenation of the GSC pool. Mutations in *hlh-30* disrupt diapause entry, maintenance and exit, causing a disconnect between nutrient cues and proper growth signaling, triggering a senescent-like phenotype. Accordingly, *hlh-30* loss induces a senescent-like DNA damage, immune and metabolic growth signature, largely reversed by reduced TGFβ signaling. In mammalian cellular models of embryonic or cancer diapause, TFEB similarly promotes survival and TFEB-TGFβ signaling is highly regulated. Of note, TFEB knockdown alone was sufficient to drastically reduce the viability of dormant cancer cells in a diapause-like state.

Our studies reveal TFEB-TGFβ axis as a primordial circuit systemically regulating diapause and stem cell quiescence/activation in response to nutrient and growth signals. Here, we showcase ARD as a tractable model to illuminate the molecular architecture of stem cell longevity, regeneration and senescence *in vivo* and developed a platform to decipher conserverd pathways governing diapause entry, maintenance and exit.

## RESULTS

### Reduced TGFβ and insulin signaling restore *hlh-30* ARD survival and progeny production

*hlh-30* ARD worms typically live no more than 5-10 days in ARD, a decrease of 88% mean life span compared to N2 wild-type (Figure 1A), while there is little effect on *ad libitum (AL)* lifespan.^16,17^ The complete collapse of *hlh-30* animals under ARD thus provided a powerful genetic selection for survivors, allowing us to dissect epistatic mechanisms acting downstream. We therefore carried out EMS mutagenesis of *hlh-30(tm1978)* mutants and screened for F2 suppressors that live at least 20 days at 20°C, and which upon refeeding, recover and reproduce (Figure 1B). We screened through 24000 genomes to isolate 15 mutants that retested true. We noticed that 8 mutants exhibited a dauer constitutive (Daf-c) phenotype under *AL* conditions at an elevated temperature of 25°C.

**Figure 1.**
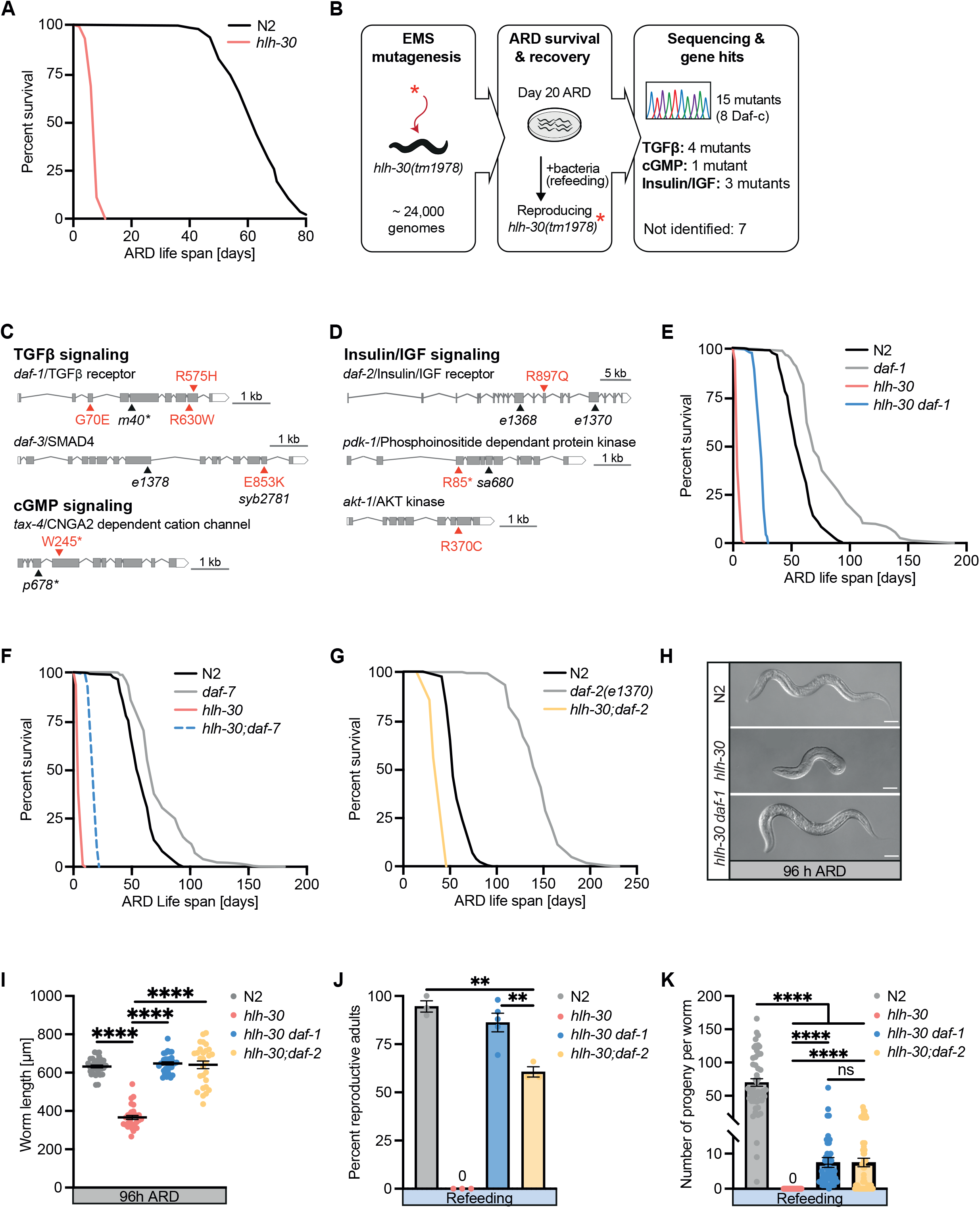
Reduced TGFβ signaling restores *hlh-30*/TFEB ARD survival and recovery. A. ARD and *AL* lifespan of *hlh-30(tm1978)* and N2 wild-type animals. B. Genetic screen for mutations that extend *hlh-30(tm1978)* ARD survival and recovery. Red star, mutation induced by EMS. C. and D. Gene structure of hit genes from TGFβ (*daf-1/TGF*β *receptor*, *daf-3/SMAD4*), cGMP (*tax-4*/*CNGA2*), and Insulin/IGF signaling (*daf-2/Insulin/IGF receptor*, *pdk-1*/*PDPK1*, and *akt-1*/*AKT*). Representative intron-exon map based on WormBase gene structures. Positions of point mutation identified in this screen are marked by red arrow heads. Amino acid alterations are given in single-letter code. Reference alleles used to validate the gene hits in black. E. and F. TGFβ signaling mutants *daf-1(m40)* (E) and *daf-7(e1372)* (F) enhance ARD survival of *hlh-30(tm1978)*. G. Insulin/IGF receptor mutation *daf-2(e1370)* enhances ARD survival of *hlh-30(tm1978)*. H. Representative images of worms at 96h of ARD. Genotypes N2, *hlh-30(tm1978), hlh-30 daf-1(m40)*. I. Body length of ARD worms at day 4. Genotypes N2, *hlh-30(tm1978), hlh-30 daf-1(m40)*, and *hlh-30;daf-2(e1370).* Each dot represents one animal. BR=3, one representative experiment. J. Percent reproductive worms refed after 10 days in ARD. Genotypes N2, *hlh-30(tm1978), hlh-30 daf-1(m40),* and *hlh-30;daf-2(e1370).* Each dot represents one experiment. K. Brood size of self-fertilizing worms refed after 10 days of ARD. Genotypes N2, *hlh-30(tm178), hlh-30 daf-1(m40),* and *hlh-30;daf-2(e1370).* Each circle represents the total progeny per worm. One representative experiment. Mann-Whitney test. BR=3. Survival curves depict one experiment. Lifespan data and statistics (log-rank tests) are presented in Table S2. ∗∗∗∗p < 0.0001; ∗∗p < 0.01. Mean ± SEM. If not stated otherwise: One-way ANOVA.

Molecular pathways governing dauer formation include TGFβ, insulin/IGF, cGMP, mTOR and steroid/bile acid-like signaling whose downregulation promote dauer and upregulation promote continuous development.^1^ Whole genome sequencing revealed that 8 mutants harbored lesions in known dauer signaling pathways, including TGFβ signaling (*daf-1*/TGFβ receptor, *daf-3*/SMAD4), insulin/IGF signaling (*pdk-1*/phosphoinositide dependent protein kinase, *akt-1*/AKT kinase, *daf-2*/Insulin/IGF receptor), and cGMP signaling (*tax-4*/cyclic nucleotide gated A2 dependent cation channel), which regulates TGFβ and Insulin-like peptide production in ciliated neurons^1,20^ (Figures 1C, 1D, marked in red), suggesting a partial overlap of dauer and ARD signaling. To validate these candidate genes, we reconstructed double mutants of *hlh-30(tm1978)* with independent reference alleles [*daf-1(m40), pdk-1(sa680), tax-4(p678), daf-2(e1368* and *e1370*)], or engineered the original mutation into the endogenous locus by CRISPR–Cas9 gene editing [*daf-3(syb2718*)] (marked in black). Consistent with the tested loci being causal, the above mutations significantly prolonged *hlh-30* ARD survival beyond 10 days, though not back to that of wild-type (Figures 1E, 1G, S1A-S1C). Notably, relative fold increase in mean life span of *hlh-30 daf-1* double mutants compared to *hlh-30* was usually greater (>3.2-4.1X) than that of *daf-1* single mutants compared to wild-type (1.1-1.2X). We also tested existent null mutants of *daf-7(e1372)*, encoding TGFβ/activin homolog, and saw similar behavior (Figure 1F). Taken together, these findings suggest that reduced TGFβ and insulin/IGF signaling act downstream or parallel to HLH-30 to promote ARD survivorship.

To consolidate our findings, we asked whether these mutations could also rescue other *hlh-30*-associated traits. Mutations in TGFβ, insulin/IGF, and cGMP signaling also rescued body size under ARD, as well as development to reproductive adult and progeny production upon refeeding (Figures 1H-1K, S1D, S1E). Notably, reduced TGFβ signaling, robustly restored the fraction of reproductive adults back to wild-type level (Figure 1J). We chose to focus mainly on TGFβ signaling for further study, since multiple hits in this pathway were found (Figures 1C and 1D), phenotypes were robust, and a role for TGFβ signaling in fasting/refeeding, quiescence/activation and longevity is less well understood than insulin/IGF signaling.

### TGF**β** signaling acts through downstream transcription factors to rescue *hlh-30* **ARD phenotypes**

To unravel how TGFβ signaling impacts *hlh-30* ARD survival, we first examined genetic epistasis interactions. With respect to dauer, DAF-7/TGFβ ligand acts through TGFβ type I and II receptors, DAF-1 and DAF-4, throughout the body to activate DAF-8 and DAF-14/r-SMADs, which inhibit DAF-3/SMAD4 and DAF-5/SNO-SKI, thereby preventing dauer and promoting continuous development (Figure 2A).^1,21^ In genetic terms, loss-of-function mutations in *daf-1*, which result in constitutive dauer formation (Daf-c) at 25°C, are suppressed by dauer formation defective (Daf-d) null mutations in *daf-3(e1376)* and *daf-5(e1386)* (Figure 2A).^22,23^ To test whether *daf-1* also genetically acts through *daf-3* or *daf-5* in ARD context, we constructed triple mutants of *hlh-30 daf-1;daf-3(e1376)* and *hlh-30 daf-1;daf-5(e1386),* using reference null alleles. As predicted, loss of *daf-3* or *daf-5* suppressed *hlh-30 daf-1* ARD longevity (Figure 2B). Additionally, *daf-3(e1376)* loss abolished *hlh-30 daf-1* progeny production, consistent with a requirement for these canonical TGFβ transcriptional mediators (Figure 2E). Downregulated TGFβ signaling also modestly promotes longevity under *AL* conditions, dependent on the DAF-16/FOXO transcription factor.^24^ We therefore constructed *hlh-30 daf-1;daf-16(mgDf50)* triple mutants, and similarly found that *hlh-30 daf-1* ARD longevity also required DAF-16(+) (Figure 2B). Last we tested *daf-12(rh61rh411)*/VDR, the major downstream transcriptional mediator of bile-acid-like steroid signaling in dauer formation^25^, but found that it was not required (Figures S2A, S2B), consistent with distinct molecular factors mediating dauer and ARD.

**Figure 2.**
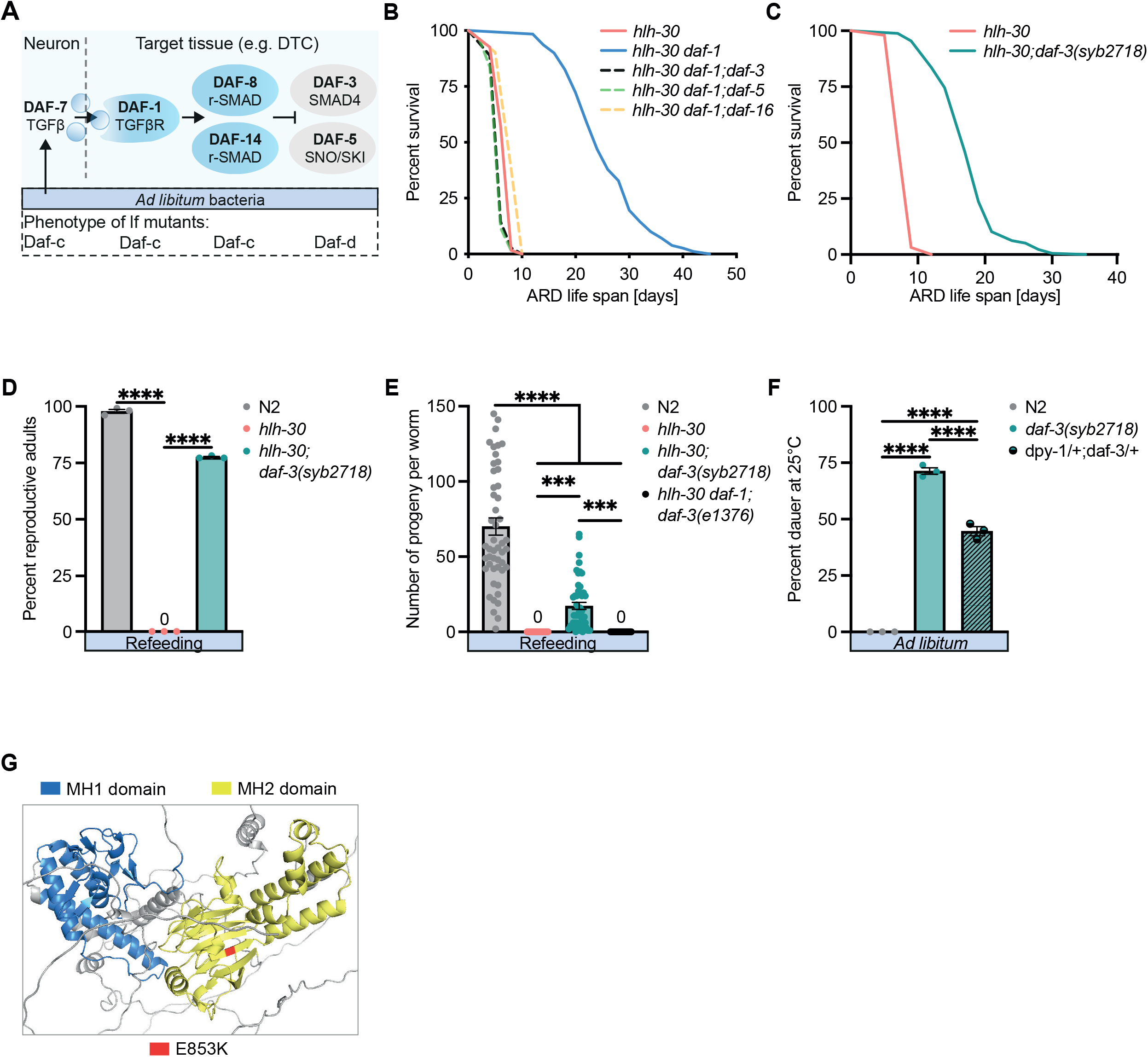
TGFβ suppression of *hlh-30* ARD survivorship and recovery depends on downstream transcription factors. A. Simplified TGFβ signaling schematic. Under *AL* conditions neuronal DAF-7/TGFβ induces a DAF-1/TGFβ receptor signal cascade inhibiting repressor activity of DAF-3/SMAD-4 and DAF-5/ SNO-SKI complex in distal tip cell (DTC) and other target tissues. Below, Daf phenotypes at 25°C of loss-of-function (lf) mutants. B. *hlh-30 daf-1* lifespan extension depends on *daf-3, daf-5* and *daf-16* transcription factors. Genotypes *hlh-30(tm1978), hlh-30 daf-1(m40);daf-3(e1376), hlh-30 daf-1;daf-5(e1386),* and *hlh-30 daf-1;daf-16(mgDf50)*. C. *daf-3*(*syb2718)* extends *hlh-30* ARD lifespan. D. *daf-3(syb2718)* promotes *hlh-30* ARD recovery and development of reproductive adults upon refeeding at day 10 ARD. Genotypes N2, *hlh-30(tm1978),* and *hlh-30;daf-3(syb2718).* Each dot represents one experiment. E. Brood size of *daf-3(syb2718)*;*hlh-30* worms refed at days 10 ARD. Genotypes as in D). Each dot represents the total progeny number of one worm. BR=3. F. *daf-3(syb2718)* exhibits a dominant Daf-c phenotype at 25°C. Genotypes N2, *daf-3(syb2718)*, and *daf-3(syb2718)/+* heterozygous. Each dot represents one experiment. G. Protein structure of *daf-3(syb2718)* showing mutation E853K in the MH2 domain in red. Conserved Mad homology 1 and 2 domains (MH1 and MH2) in blue and yellow, respectively. ∗∗∗∗p < 0.0001; ∗∗∗p<0.001. Mean ± SEM (D-F). One-way ANOVA.

Of note, the *daf-3(syb2718)* allele obtained from our screen behaved exactly opposite to the *daf-3* null allele, suggesting *syb2718* is a gain-of-function. Indeed, *daf-3(syb2718)* mutants exhibited Daf-c phenotypes at 25°C (*AL*), and potently restored *hlh-30* ARD survival, recovery and reproduction (Figures 2C-2F). Consistent with gain-of-function, *daf-3(syb2718)*/+ heterozygotes were dominant for Daf-c phenotypes (Figure 2F). Further, the changed amino acid (E853K) reverses charge, falls within the MH2 domain (Figure 2G), and is predicted to disrupt inhibitory interactions with r-SMADs, uncoupling DAF-3/SMAD4 activity from upstream inputs.^26^ Altogether these findings reveal that reduced TGFβ signaling rescues *hlh-30* survivorship and recovery, and point to DAF-3/SMAD4 as a key downstream transcription factor promoting protective mechanisms within these circuits.

### HLH-30/TFEB regulates TGF**β** signaling in ARD

The genetic epistasis experiments described above suggest that HLH-30(+) could act to downregulate TGFβ signaling under ARD. To directly address this idea, we first measured expression of *daf-7/*TGFβ ligand, using a *daf-7p::gfp* reporter that expresses *gfp* under the *daf-7* promoter. Under fed conditions *daf-7p::gfp* is expressed mostly in the ASI and ASJ amphid sensory neurons.^21^ These neurons secrete TGFβ, serving as an endocrine source to regulate TGFβ receptor signaling in target tissues throughout the body.^21,27^ In wild-type, *daf-7p::gfp* expression in ASI was low during ARD, but high upon refeeding/recovery (Figures 3A-3C). In *hlh-30* mutants, by contrast, *daf-7p::gfp* expression in ASI was high in ARD, but not further upregulated upon refeeding/recovery, suggesting that HLH-30(+) normally inhibits *daf-7* expression in ARD and possibly induces it during recovery. Endogenously tagged *hlh-30::mNeonGreen* itself was expressed in the ASI, rapidly entered the nucleus within 2 hours of ARD induction and persisted there during early ARD (Figures 3D, 3E). Upon refeeding, HLH-30::mNeonGreen promptly exited the nucleus, suggesting that HLH-30 directly or indirectly inhibits *daf-7* expression cell-autonomously in ASI in response to nutrient cues. In addition to *daf-7p::gfp* expression in ASI, we also observed a low level *daf-7p::gfp* expression in neurons anterior to the nerve ring (possibly OLQ or IL1, based on CENGEN data, https://cengen.shinyapps.io/CengenApp/), revealing a novel expression pattern under ARD (Figures 3F, S3A). *hlh-30* mutants also perturbed this regulation, in this case causing lower *daf-7p::gfp* expression at 96 hours of ARD compared to wild-type. Nevertheless, overall neural *daf-7p::gfp* expression was still higher under ARD in *hlh-30* compared to wild-type.

**Figure 3.**
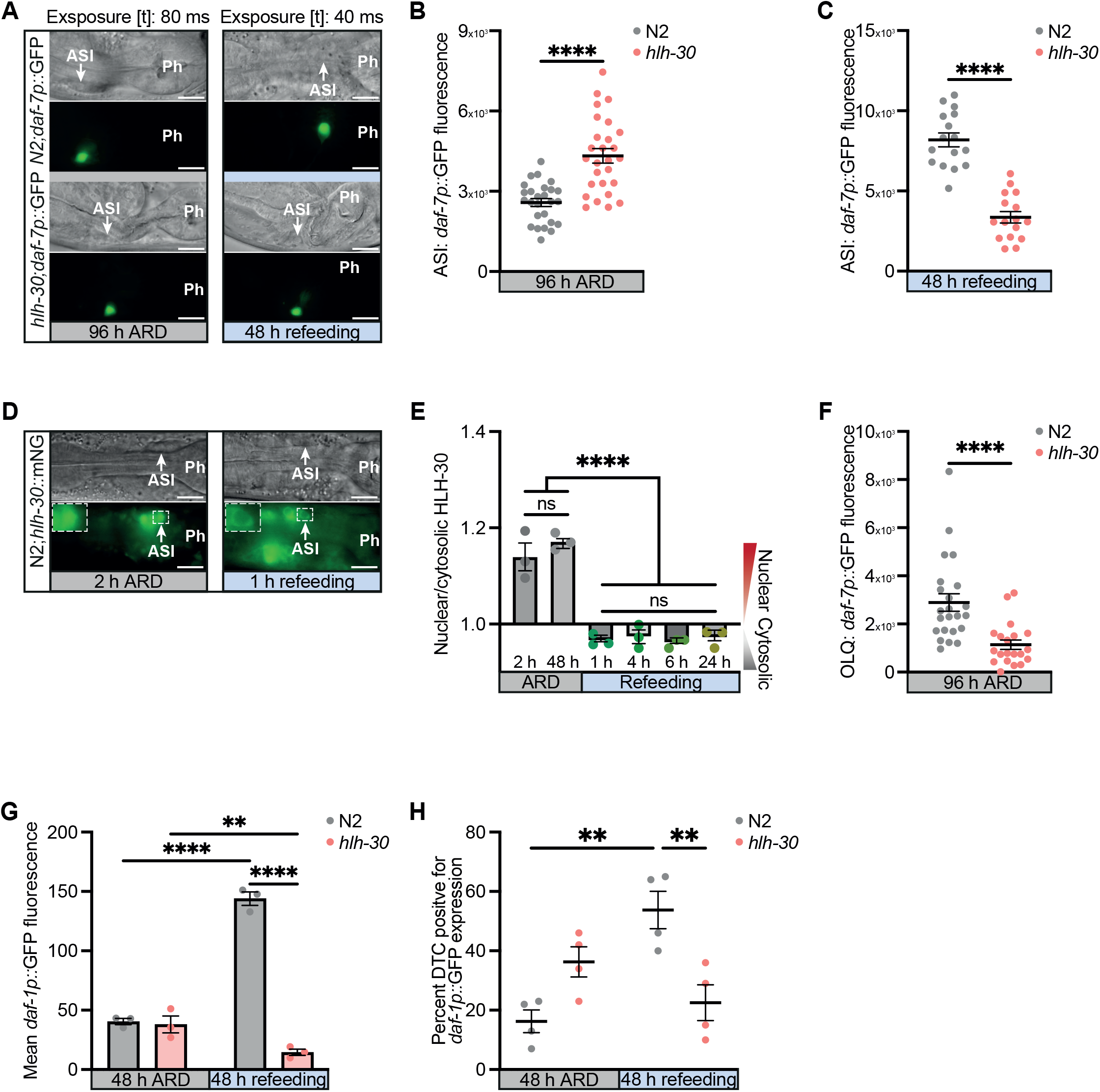
HLH-30/TFEB regulates TGFβ signaling. A. *daf-7p::gfp* expression (TGFβ) in ASI neurons of N2 and *hlh-30(tm1978)* at 96 hours of ARD and after 48 hours of refeeding. DIC and fluorescent images of head region of N2 and *hlh-30*. Arrows pointing to ASI. Ph, terminal bulb. Fluorescent images exposure time: ARD 80 ms. Refeeding 40 ms. BR=3. B. and C. Quantification of *daf-7p*::*gfp* expression in ASI at 96 hours of ARD (B, 80 ms exposure time) and after 48 hours of refeeding (C, 40 ms exposure time). Each dot represents the *daf-7p::gfp* expression of one ASI neuron. BR=3, one representative experiment shown. D. *hlh-30::mNeonGreen* (*hlh-30::*mNG) expression pattern in ASI neurons at 2 hours of ARD and upon 1 hour of refeeding 48 hours ARD worms. DIC and fluorescent images of head region of N2;*hlh-30::*mNG. Arrow pointing to ASI. Ph, terminal bulb. BR=3. E. Nuclear/cytosolic *hlh-30::*mNG expression in ASI under ARD (2 and 48 hours) and recovery (1, 4, 6 and 24 hours of refeeding 2 ARD worms). Each dot represents one experiment. BR=3. F. Quantification of *daf-7p::GFP* expression in OLQ neurons of N2 and *hlh-30(tm1078)* at 96 hours of ARD. Exposure time 400 ms. Each dot represents the *daf-7p::*GFP expression in one OLQ neuron. BR=3, one representative experiment. G. Quantification of whole body (WB) *daf-1p*::GFP (TGFβ Rec Type 1) at 48 hours of ARD and 48h of refeeding measured with COPAS biosorter. BR=2 one representative experiment. H. Percent DTC positive for *daf-1p::*GFP expression of N2 and *hlh-30(tm1978)* at 48 hours of ARD and after 48 hours refeeding. Scale bar 10 μm. ∗∗∗∗p < 0.0001; ∗∗∗p<0.001; ∗∗p<0.01; ∗p<0.05; ns, no significant difference. Mean ± SEM. Mann Whitney test (B, C, F). One-way ANOVA (E). Two-way Anova (G).

We next examined regulation of *daf-1p::* GFP, using a Copas biosorter to obtain unbiased whole body fluorescent analysis of hundreds of animals. While expression levels of *daf-1p::*GFP were not significantly different in ARD, *hlh-30* animals consistently expressed lower levels of *daf-1p::gfp* upon refeeding and were unable to adapt to food cues compared to wild-type (Figure 3G). To examine tissue-specific regulation, we then monitored *daf-1p::*GFP by microscopy within the distal tip cells (DTC), which comprise the niche directly regulating GSC proliferation and differentiation. In line with our *daf-7/TGF*β and whole body *daf-1* data wild-type, *daf-1p::*GFP expression was low in the DTC under ARD, but high upon refeeding (Figure 3H). In *hlh-30* mutants by contrast, mean *daf-1p::*GFP expression was increased by 27% compared to wild-type during ARD, and did not respond to refeeding, showing a misregulation of *daf-1* receptor expression in response to nutrient cues.

In sum, HLH-30(+) downregulates TGFβ signaling during ARD via inhibition of *daf-7*/TGFβ in ASI and *daf-1*/TGFβ receptor in the GSC niche, and this regulation is reversed upon recovery. Nutrient regulation of TFEB-TGFβ signaling from neuron to reproductive system suggest an integral role of this axis in controlling quiescence and activation via a systemic mechanism.

### HLH-30 regulates GSC dynamics via TGF**β** and Notch signaling

The gonad encompasses the only known stem cell compartment in the adult worm. It is capped by the DTC, which serves as the niche regulating germline stem activation and spatially organizes the mitosis/meiosis transition (Figure 4A). This spatial organization relies in part on the DTC extending long processes which expand the zone of stem cell activation.^28,29^ Both germline morphology and signaling are altered under ARD.^15^ In response to nutrient deprivation, the gonad retracts and GSCs become quiescent and cease proliferation. Upon refeeding, GSCs exit dormancy, resume proliferation and differentiate, culminating in progeny production even after prolonged starvation.^16,30^

**Figure 4.**
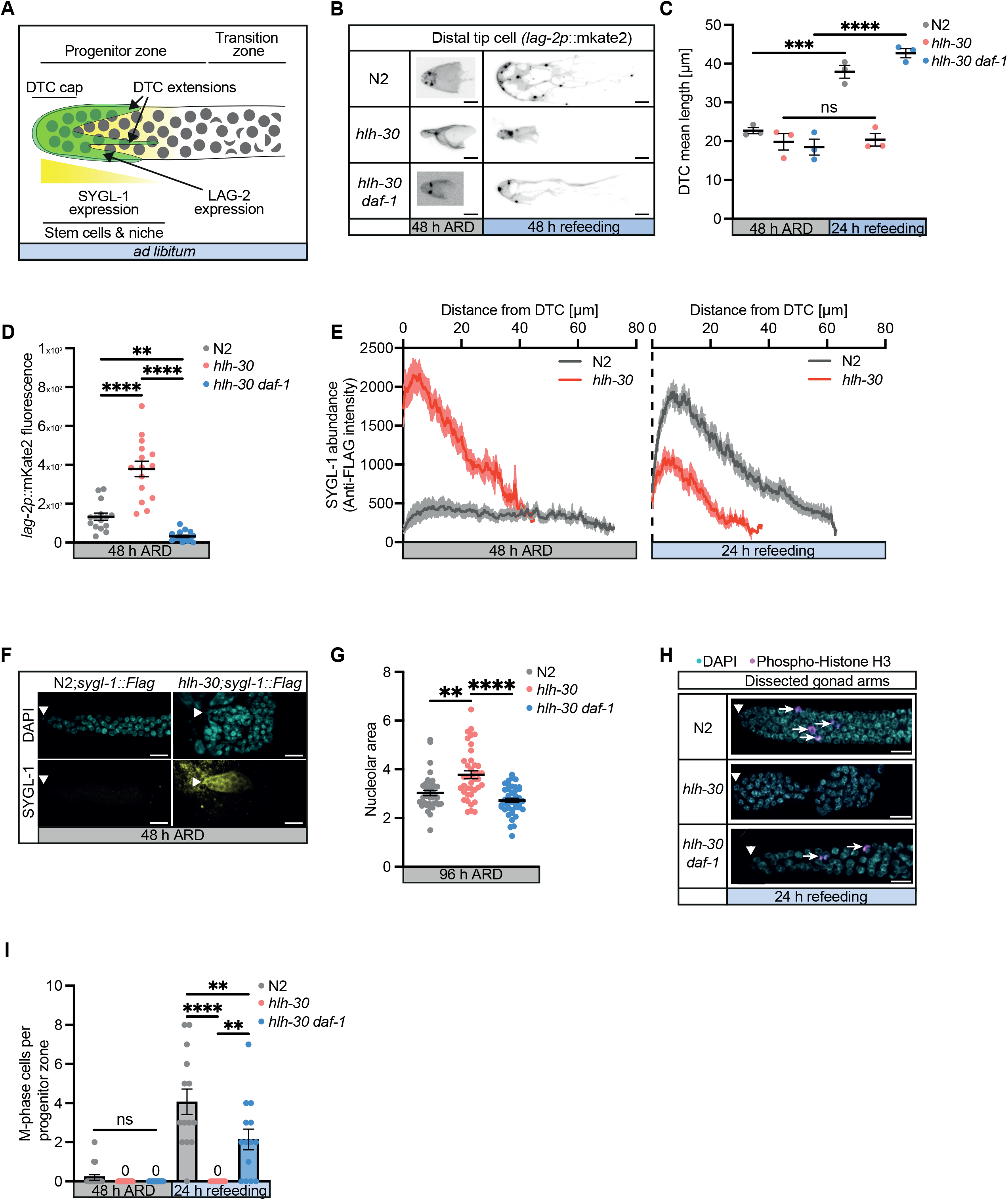
Regulation of Notch signaling and germ cell phenotypes by *hlh-30* and *daf-1*. A. Hermaphrodite distal gonad schematic. Distal tip cell (DTC) cap and extensions green. Cell nuclei in gray. The progenitor zone contains a distal pool of stem cells (area of SYGL-1 expression, yellow) and a proximal pool of cells that are differentiating. The transition zone contains crescent-shaped nuclei characteristic of early meiotic prophase. B. *lag-2p::mKate2::PH* (DSL ligand) expression in DTC at 48 hours of ARD and after 48 hours refeeding. Fluorescent pictures. Contrast and intensity of the pictures were adjusted to visualize processes extending from the cap structure. Genotypes N2, *hlh-30(tm1978), hlh-30 daf-1(m40)*. BR=3. C. DTC extension measured from the cap to the end of the most distal process. *lag-2p::mKate2::PH*. Each dot represents one experiment. One-way ANOVA. D. Quantification of *lag-2p::mKate2::PH* expression intensity in the DTC cap at 48 hours of ARD. Each dot represents one experiment. E. Quantification of SYGL-1 (Notch target) abundance in the distal gonad of 48-hour-ARD-worms and of 24-hour refed worms, based on intensity of α-FLAG staining. *sygl-1::3xFlag* in N2 and *hlh-30(tm1978)* background. Average intensity values (Y-axis) were plotted against distance (μm) from the DTC (X-axis). Lines, mean intensity; shaded areas, SEM; BR=3. F. Representative images of SYGL-1 abundance stained with DAPI (DNA, turquoise) and anti-3xFLAG (SYGL-1, yellow). 10 μm, Arrowhead, DTC. G. Quantification of germ cell nucleolar area of N2, *hlh-30(tm1078), hlh-30 daf-1(m40)* at 96 hours of ARD. BR=3. One representative experiment. H. Distal gonad arms dissected from worms, refed at 48 hours of ARD for 24 hours, stained with DAPI (DNA, turquoise) and anti-phospho-histone H3 (M-phase chromosomes, magenta). Genotypes N2, *hlh-30(tm1978), hlh-30 daf-1(m40)*. Scale bar 10 μm, Arrowhead, DTC. I. Number of M-phase cells in the distal gonad arms (progenitor zone) of N2, *hlh-30*, *hlh-30 daf-1* at 48 hours ARD and 24 hours refeeding. BR=3., one representative experiment. ∗∗∗∗p < 0.0001; ∗∗∗p<0.001; ∗∗p<0.01; ns, no significant difference. Mean ± SEM. One-way ANOVA.

GSCs are primarily regulated via Notch signaling.^31^ Under *AL* conditions, the LAG-2/Delta-Serate ligand expressed in the DTC niche binds to the GLP-1/Notch receptor in adjacent GSCs, to promote expression of target genes including *sygl-1,* resulting in stem cell activation and mitosis, and prevention of premature meiosis.^32,33^ Given that TFEB-TGFβ impacts progeny production (Figures 1K, 2E), we wondered whether it correspondingly affects Notch signaling and GSC function during ARD and recovery. We measured three parameters: *lag-2p::mKate2::PH* expression level in the DTC (niche), DTC long process extension (niche remodeling, germ cell zone of activation), and *sygl-1::3xFlag* abundance (GSCs) as proxy measures of Notch activity. In wild-type, we observed that all three parameters were reduced under ARD, and DTC extension and *sygl-1::3xFlag* expression increased upon recovery, consistent with downregulation of Notch signaling in quiescence and upregulation during activation, respectively (Figures 4B-4F, S4A). By contrast, *hlh-30* mutants exhibited enhanced *lag-2p::mKate2::PH* and *sygl-1::3xFlag* expression under ARD compared to N2, yet during recovery failed to extend LAG-2 DTC processes, and limited *sygl-1::3xFlag* expression, suggesting *hlh-30* loss dysregulates Notch signaling during both quiescence and activation in both niche and stem cells. As predicted, *daf-1* mutation reversed many features of this *hlh-30* dysregulation (Figures 4C, 4D).

We hypothesized that *hlh-30* mutant GSCs might not successfully enter stem cell quiescence during ARD fasting due to upstream pro-growth signals. Therefore, we examined GSC morphology by DIC optics and expression of the mitotic marker phospho-Histone3 (H3) by fluorescent microscopy (Figures 4G-4I, S4B). In wild-type animals in ARD, GSC nucleoli appeared compact and nuclei did not express phospho-H3, consistent with quiescence. During recovery, germ cell nuclei expressed phospho-H3, consistent with quiescence exit and cell proliferation. By comparison, *hlh-30* mutants in ARD exhibited enlarged germ cell nuclei^16^ and nucleoli (Figure 4G, S4B). Upon refeeding, *hlh-30* germ cells failed to express phospho-H3 and were unable proliferate (Figures 4H, 4I). *daf-1* mutation restored small nucleoli to *hlh-30* mutants during ARD, and phospho-H3 expression during recovery. These observations are consistent with the idea that *hlh-30* mutants fail to exert proper cell cycle control in response to fasting/refeeding, and that germ cell function and dynamics can be restored by inhibition of TGFβ signaling.

### *daf-1* mutation enables fasting adaptive transcriptome in *hlh-30* mutants

To gain further insight into mechanism, we sought to identify differentially regulated genes (DEGs) and processes during ARD and recovery. We performed RNA-seq analysis with N2, *hlh-30(tm1978), daf-1(m40)* and *hlh-30 daf-1* double mutants at 48 hours ARD and 12 hours of refeeding (REF) to capture relatively early differences in gene expression. We chose 48 hours to ensure full ARD maturation, yet minimize changes associated with *hlh-30*-induced morphological alterations and early death. Principle component analysis revealed clear separation of all genotypes under ARD and refeeding (Figure 5A), showing good data quality. Hierarchical clustered expression heat maps showed segregation of *hlh-30* from all other genotypes, clustering under both conditions more closely to ARD, while *hlh-30 daf-1* clustered more closely to N2, suggestive of dysregulation of gene expression in *hlh-30* and reversion by *daf-1* in both states (Figure 5B).

**Figure 5.**
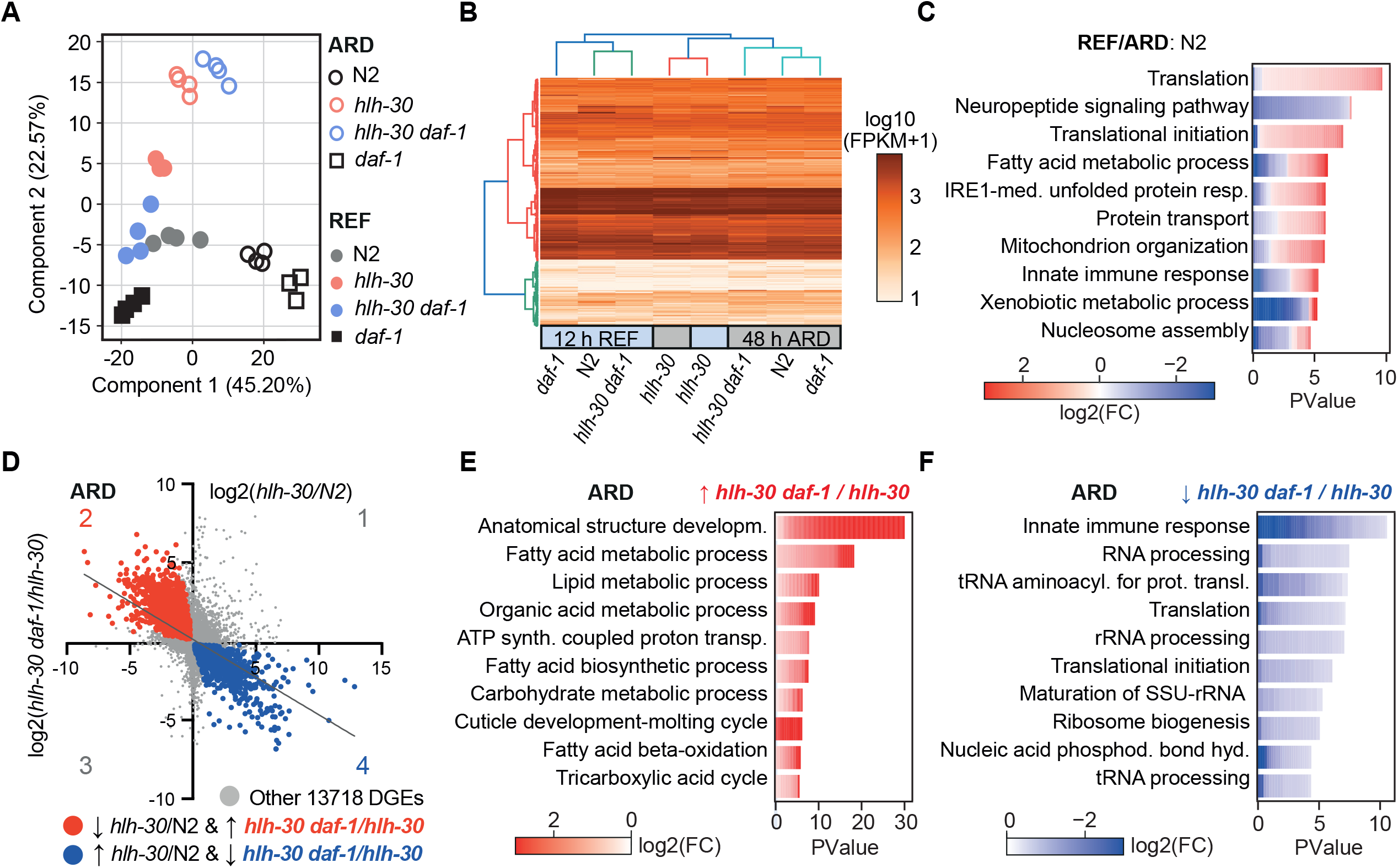
Transcriptomic analyses of *hlh-30* and *daf-1* interactions. **A.** Principal component analysis of RNA-Seq data of N2, *hlh-30(tm1978), daf-1(m40),* and *hlh-30 daf-1* at 48 h ARD and after 12 h refeeding/recovery (REF). Each dot represents one sample. **B.** Heatmap of protein coding genes with padj < 0.05 (RNA seq data) of indicated genotypes under 48 hours of ARD (gray box) and 12 hours REF (blue box). The dendrograms show hierarchical clustering by Euclidean distance. **C.** Gene ontology enrichment analysis of 9169 DEGs (padj < 0.05) in N2 REF. Top 10 biological processes. Complete GO term list is shown in Table S3. **D.** Correlation plot of *hlh-30(tm1978)*/N2 and *hlh-30 daf-1(m40)*/*hlh-30* genes at 48 hours of ARD. Significant regulated genes (padj < 0.05) highlighted in red (genes down in *hlh-30* and reversed by *daf-1,* 2347 DEGs, quadrant 2) or blue (genes up in *hlh-30* and reversed by *daf-1*, 2083 DEGs, quadrant 4). Other 13718 genes, gray. Simple linear regression line in gray. Equilibration Y = - 0.48X + 0.13; R^2^ = 0,27. **E.** Gene ontology enrichment analysis of 2347 DEGs down in *hlh-30(tm1978)/*N2 ARD and up in *hlh-30 daf-1(m40)/hlh-30* ARD (padj < 0.05, quadrant 2). Top 10 biological processes. Complete GO term list is shown in Table S3. **F.** Gene ontology enrichment analysis of 2083 DEGs up in *hlh-30(tm1978)/*N2 ARD and down in *hlh-30 daf-1(m40)/hlh-30* ARD (quadrant 4). Top 10 biological processes. Complete GO term list is shown in Table S3.

To dissect expression changes during ARD and upon refeeding, we first compared wild-type transcriptomes (REF/ARD), resulting in approximately 9000 DEGs (padj. p<0.05). Gene ontology enrichment revealed biological processes (BPs) like translation, unfolded protein response, fatty acid metabolism and mitochondrial organization mostly upregulated and neuropeptide signaling and innate immune response downregulated (Figure 5C). By inference, perception and integration of starvation information result in metabolic adaption towards survival and catabolism. Refeeding N2 induces proteolytic turnover and anabolic processes linking nutrient supply with growth and reproduction.

Three key processes differentially regulated during N2 REF/ARD, namely translation, fatty acid metabolism and innate immune response, were dysregulated in *hlh-30* mutants, both during ARD and upon refeeding (*hlh-30*/N2, Figures S5A, S5B), confirming the major role of HLH-30(+). This observation led us to hypothesize that *hlh-30* mutants fail to properly respond to ARD nutrient restriction and refeeding. Unable to adapt proteostasis, remodel essential metabolic and growth processes, *hlh-30* worms might enter a senescent-like state with dysregulated immune signature during ARD.

Next, we focused on processes associated with *hlh-30* phenotypic rescue by *daf-1.* Reasoning that genes downregulated in *hlh-30/*N2 might be upregulated in *hlh-30 daf-1*/*hlh-30* and vice versa, we found a large fraction of such DEGs negatively correlated, consistent with a profound phenotypic reversal under ARD and upon refeeding (Figures 5D, S5C). ARD DEGs down in *hlh-30/*N2 and reversed by *daf-1* were mainly enriched in metabolic processes, *e.g.,* fatty acid beta-oxidation and TCA cycle (quadrant 2, 2434 DEGs, Figures 5D, 5E). Refeeding revealed downregulation of translation, fatty acid and mitochondrial metabolism, heterochromatin and other processes in *hlh-30*/N2, reversed by *daf-1*(1500 DEGs, quadrant 2, Figures S5C, S5D).

ARD DEGs up in *hlh-30/*N2 and reversed by *daf-1* included innate immune response, translation, and nucleic acid phosphodiester bond hydrolysis (quadrant 4, 2083 DEGs, Figures 5D, 5F). Refeeding DEGs reversed by *daf-1* were associated with innate immune response, nucleic acid phosphodiester bond hydrolysis, PERK-mediated UPR, response to gamma radiation, and neuropeptide signaling pathway (2173 DEGs, quadrant 4, Figure S5C, S5E). In conclusion, *daf-1* restored dysregulated immune and growth metabolic signatures of *hlh-30* mutants, enabling them to adapt to ARD or to recover upon refeeding.

Our studies above revealed DAF-3/SMAD4 as a key mediator of *hlh-30 daf-1* survivorship and reproduction. SMAD4 loss disrupts DNA repair mechanisms leading to enhanced genomic instability and inflammation in mice and human cancer.^34,35^ The GO term “DNA repair” was enriched among HLH-30 targets, as seen in published HLH-30 ChIP-seq data under ARD.^16^ Here, we also found GO terms DNA damage (ARD), repair (ARD and REF) and related terms such as phosphodiester bond hydrolysis (ARD and REF), response to gamma radiation (REF), and apoptotic DNA fragmentation (REF) dysregulated in *hlh-30* and reversed by *daf-1*. Altogether these findings suggest that *hlh-30* mutation causes genomic instability, dysregulated immune response and a senescent-like state.

In line with our previous findings (figure 3) the transcription factors *daf-3/SMAD4* and *hlh-30/TFEB* showed similar expression trends in worms, being upregulated in N2 ARD compared to refeeding (Figure 7A). We recently demonstrated that the fasting/refeeding response is dysregulated in aged short-lived killifish (*Nothobranchius furzeri*) (Ripa, accepted), and thus sought to evaluate the expression of *tfeb* and *smad4* in this model organism. Of note, similar to the worm, killifish *smad4* and *tfeb* were upregulated during fasting and low upon refeeding in different tissues (Figure 7B). Furthermore, their regulation by fasting/refeeding appeared compromised with aging, consistent with our previous results (Ripa, accepted). These observations hint at conservation of a TFEB-SMAD4 axis in vertebrates and underscore the potential significance of this axis in nutrient sensing, metabolic flexibility and the aging process.

### *hlh-30* mutants exhibit senescent-like phenotypes

As described above, *hlh-30* mutants exhibit an aberrant form of quiescence in which animals are unable to recover upon refeeding. Mutants show a transcriptional signature of dysregulated growth and metabolism, innate immune signaling and DNA damage, as well as elevated TGFβ and Notch signaling, and enlarged germ cells that fail to enter mitosis upon refeeding. We realized these phenotypes are strikingly reminiscent of cellular senescence (Table S1).

The study of cellular senescence has been little addressed in the worm, despite its use as an organismal aging model. To further explore possible features of senescence in *hlh-30* mutants, we first asked whether global changes in the transcriptome could give hints. The biological age of worms can be predicted by binarized transcriptomic aging (BiT age) analysis.^36^ Using this transcriptome clock, we observed *hlh-30* ARD worms to be significantly older than N2 and *hlh-30 daf-1* mutants in ARD (Figure 6A). Accordingly, *hlh-30* ARD worms showed a decline in measures of healthspan, namely body movement and pharyngeal pumping, already at 48 hours of ARD; such behavioral phenotypes were not seen in wild-type at day 10 ARD^16^, suggesting premature physiological aging in *hlh-30* (Figures 6B, 6C).

**Figure 6.**
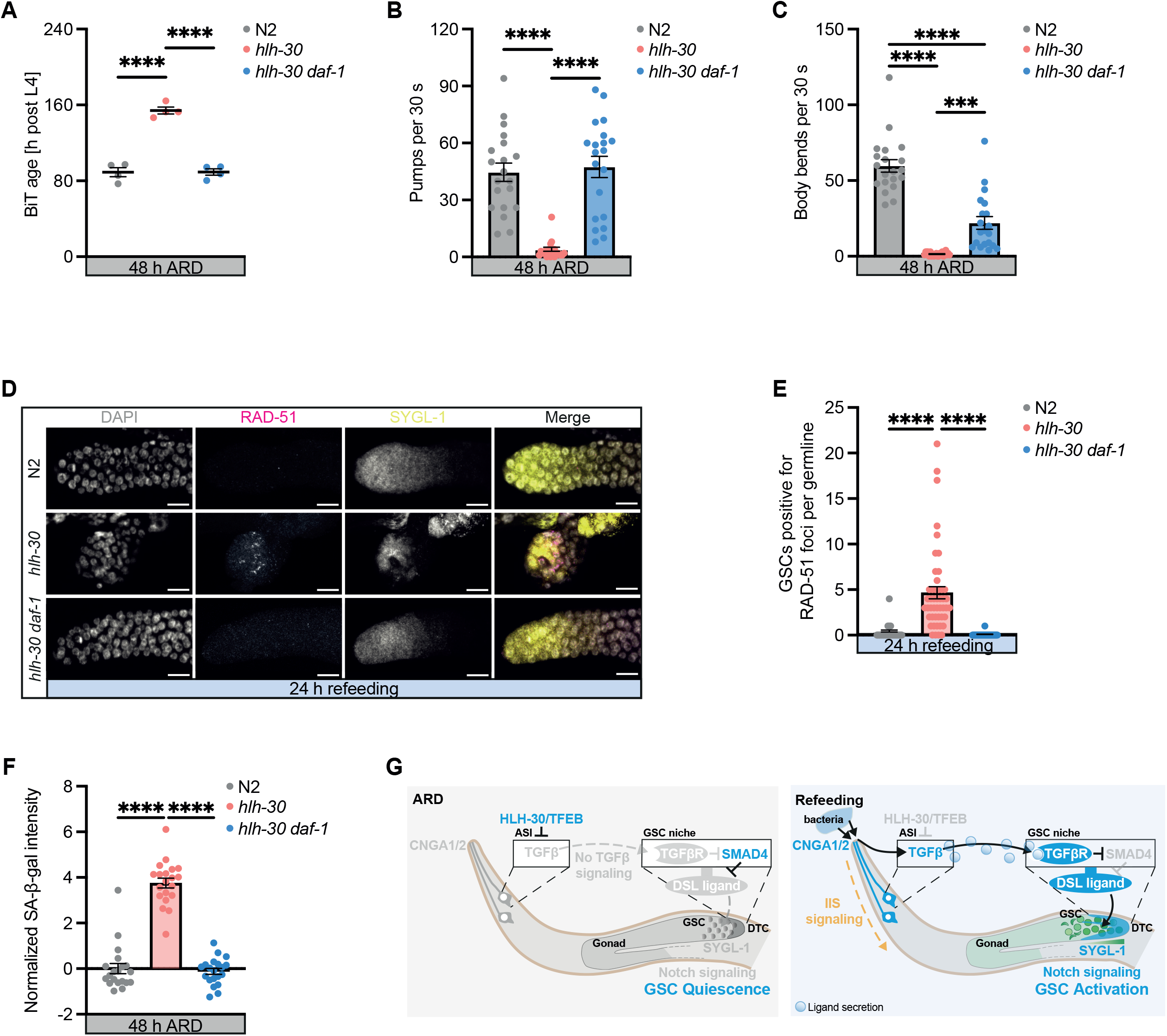
*hlh-30* mutants exhibit senescent-like phenotypes. A. Biological age prediction (BiT) from transcriptomes of N2, *hlh-30(tm1978)*, and *hlh-30 daf-1(m40)* worms at 48 hours of ARD. Each point represents one replicate. B. and C. Pumping (B) and Body bending rates (C) of 48 hours ARD worms. N2, *hlh-30(tm1978), hlh-30 daf-1(m40)*. 30 second intervals. Each dot represents the pumping rate or body bends of one animal. BR=3, one representative experiment. D. Photomicrographs of distal gonad arms of *sygl-1::3xFlag* worms in N2, *hlh-30(tm1978), hlh-30 daf-1(m40)* backgrounds, stained with DAPI (nucleus) anti-3x FLAG (GSCs zone) and anti-RAD-51 (DNA damage foci) antibodies. Scale bar 10 μm. E. Quantification of GSCs positive for RAD-51 foci in the *sygl-1*-positve area per gonad. One dot represents the number of positive GSC per gonad arm per worm. Pooled from 3 independent BRs. F. SA-β-gal RGB color intensity in the head region of 48 h ARD worms, normalized to background. Genotypes N2, *hlh-30(tm1978), hlh-30 daf-1(m40)*. Each dot represents the SA-β-gal intensity per worm head. BR=3. one representative experiment. G. TGFβ-TFEB working model. In wild-type ARD HLH-30/TFEB is nuclear localized (active) and downregulates DAF-7/TGFβ ligand in ASI neurons. When TGFβ signaling is low, DAF-3/SMAD4 is active and inhibits Notch signaling via LAG-2/DSL, and GSCs are quiescent. Under refeeding, neuronal HLH-30 exits the nucleus, TGFβ signaling is active, and DAF-1/TGFβ inhibits DAF-3/SMAD4. Hence, Notch signaling is active, stem cell niche remodeling (DTC outgrowth) is initiated, GSCs are activated and worms reproduce. cGMP and IIS work upstream or in parallel to TGFβ signaling. Details see text. ∗∗∗∗p < 0.0001; ∗∗∗p<0.001. Mean ± SEM (A-C, E, F). One-way ANOVA.

In response to severe DNA damage, cells cease division and can become senescent, and remain unable to divide.^37^ Since *hlh-30* mutants showed an elevated DNA damage response in transcriptome data (above), we wondered whether molecular correlates of DNA damage were visible. For these experiments, we stained the extruded gonads of animals with anti-RAD-51 antibodies to visualize DNA damage foci in GSCs.^38^ In control N2 animals maintained under *AL* conditions and exposed to ionizing radiation, DNA damage foci were readily apparent in GSCs, while in N2 animals experiencing ARD recovery without radiation, no such foci were visible (Figures 6D, 6E, S6A). By contrast, in *hlh-30* mutants we observed an elevated fraction of animals harboring germ cells with RAD-51 foci under conditions of ARD recovery, suggesting *hlh-30* mutants experience DNA damage or have less efficient DNA repair when attempting to resume growth, which is reversed by *daf-1* mutation (Figures 6D, 6E).

Many senescent cells also express senescence associated β-galactosidase (SA-β-gal) which is thought to reflect dysregulated lysosomal function. Previous studies show that worms exhibit elevated endogenous SA-β-gal activity as they age, which is accelerated upon exposure to high salt^39^, a stress that induces DNA breaks and senescence in cell culture. Interestingly, we found that *hlh-30* worms in ARD exhibited a large increase in SA-β-gal staining compared to wild-type controls. Further, *daf-1* mutation suppressed this phenotype (Figure 6F, S6B).

Dysregulation of autophagy affects tissue homeostasis and accelerates cellular senescence.^40^ TFEB is considered a major regulator of autophagy, which made us wonder whether *hlh-30* collapse results from autophagy failure. Therefore, we tested autophagy mutants *atg-7*/*ATG7* (early autophagy) and *lgg-2/ATG8*, (phagosome maturation and degradation). Surprisingly, we saw no significant difference in *atg-7* brood size upon recovery from ARD day 10 in comparison to wild-type (Figure S6C). *lgg-2* mutants showed a mild brood size reduction of 36%, however under *AL* conditions we also observed a reduction in total brood size (Figures S6D & S6E). Together these results indicating that other HLH-30/TFEB target processes are more important to ARD survivorship and recovery.

Altogether these findings are consistent with the idea that ARD induces an aberrant cellular and organismal quiescence in *hlh-30* mutants that resembles cellular senescence, reversed by reduced TGFβ signaling. Based on our observations we propose the following working model (Figure 6G). In response to ARD fasting, HLH-30(+) normally downregulates TGFβ within ASI sensory neurons and TGFβ type 1 receptor in the DTC stem cell niche, resulting in lower systemic TGFβ signaling and activation of DAF-3/SMAD4. Consequently, *lag-2*/DSL is downregulated in the GSC niche, and *sygl-1* in the GSCs, resulting in reduced Notch signaling, and reproductive quiescence. Upon refeeding, TGFβ and Notch signaling are induced, resulting in stem cell activation and reproductive competence. In *hlh-30* mutants, there is a mismatch between nutrient cues and growth signaling regulation. Consequently, TGFβ and Notch signaling are improperly derepressed in diapause, triggering germline dysfunction, while during refeeding, these signaling pathways are not induced, stem cells become senescent and don’t recover.

### TFEB plays an essential role in mammalian diapause models

Having found preliminary evidence for a conserved TFEB-SMAD4 axis in vertebrates from our RNA seq data (Figures 7A and 7B), we asked if the role of TFEB was conserved in other diapause models, such as mouse embryonic stem cells (mESCs) diapause^41,42^ and in the recently reported human cancer cell diapause-like state.^7,8^ In both cases, the state of diapause (or diapause-like) can be mimicked *in vitro* using INK128 (also known as Sapanisertib) a dual inhibitor of mTOR/PI3K.^7,42^ From the analysis of a genome-wide CRISPR-Cas9 screen performed in mESCs, we identified TFEB as a major determinant of diapause survival (Ramponi et al, in preparation). In particular, a doxycycline-inducible Cas9 was activated in either proliferating control mESCs or in INK128-diapause mESCs, both carrying a genome-wide library of short-guide RNAs (sgRNAs).^43^ After deep sequencing the abundance of the different sgRNAs was determined. Activation of Cas9 in control mESCs did not alter the abundance of sgRNAs for TFEB (Figure 7C). In contrast, activation of Cas9 resulted in a remarkable loss of sgRNAs for TFEB in diapause mESCs. This indicates that TFEB is essential for the survival of mESCs in diapause, but not for proliferating mESCs. Similarly, *hlh-30* mutation in *C. elegans* shows little effect on wild-type *AL* survival but is essential for ARD.^16,17^

**Figure 7.**
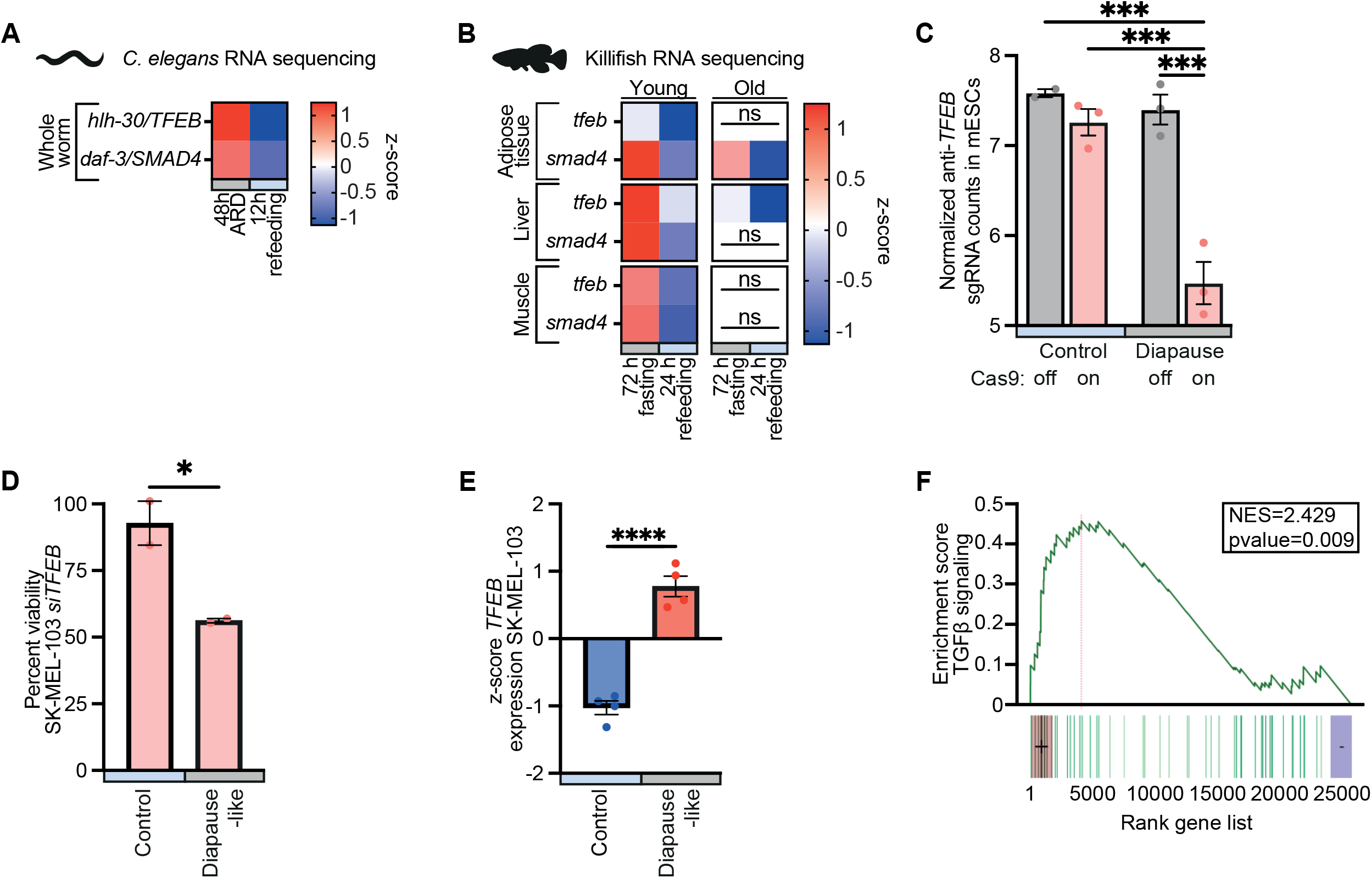
TFEB-TGFβ axis plays an essential role in fasting/refeeding and diapause across species. A. Heat map of *C. elegans hlh-30* and *daf-3* expression from RNA seq data of wild-type at 48 hours of ARD and after 12 hours of refeeding/recovery. Z score. BR=4. B. Heat map of killifish *tfeb* and *smad4* expression from adipose, liver and muscle tissues of young and old fasted and refed fish. RNA seq samples. Z score. BR=4. C. sgRNAs counts for TFEB in the four analyzed conditions: control proliferating mESCs, control proliferating mESCs with doxycycline to induce the Cas9 activity, mESCs in diapause (by treatment with INK128) and mESCs in diapause treated with doxycycline. BR=3. Two-way ANOVA D. Effect of TFEB silencing on the viability of Sk-Mel-103 cells, when proliferating and in the diapause-like state. Diapause-like state was induced by treatment with INK128. Each column is normalized on its respective siRNA scrambled. BR=2. Unpaired t test. E. TFEB expression levels from RNAseq datasets of proliferating compared to diapause-like Sk-Mel-103 cells. BR=4. Unpaired t test. F. GSEA enrichment plot of the TGFβ signaling pathway genes in diapause-like Sk-Mel-103 compared to proliferating Sk-Mel-103.

We wondered if the role of TFEB in embryonic stem cells could be extended to diapause-like human cancer cells. For this, we used human melanoma SK-MEL-103 cells that underwent a robust and reversible diapause-like arrest in the presence of INK128 (Ramponi et al, in preparation). Interestingly, siRNA-mediated downregulation of TFEB was able to reduce the survival of human melanoma diapause-like cells by almost 50%, while it had no effect on proliferating cancer cells (Figure 7D). RNAseq data from the same cell line in diapause indicated that there is an upregulation in the levels of TFEB mRNA during diapause (Figure 7E), which further supports an active role of TFEB in the survival of these cells. Gene Set Enrichment Analysis (GSEA) also indicated that the diapause-like state in human cancer cells is accompanied by an altered TGFβ signaling (Figure 7F). It is important to note that, in mammalian cells, TGFβ signaling inhibits proliferation^44^, while the same pathway in worms promotes proliferation. Therefore, in both systems, proliferation is inhibited during diapause: in mammalian cells by activating TGFβ, and in worms by inhibiting TGFβ.

## Discussion

In this work, we postulated that the study of the worm adult reproductive diapause can illuminate primordial molecular mechanisms underlying quiescence and activation in the adult animal, and yield insights into regeneration and longevity from the cellular to organismal levels. Focusing on ARD allowed us to precisely monitor germ stem cell dynamics *in vivo* and thus pinpoint regulatory pathways governing the quiescence and revival of the sole stem cell pool in the adult worm. Notably, the balance between quiescence and activation must be tightly controlled, since irreversible quiescence can lead to regenerative failure, while overactivation can deplete the stem cell pool, impacting disease, cancer and senescence.^45–47^ It follows that how stem cells and their niches maintain this cellular plasticity throughout life is critical to healthy aging. Additionally, fasting and refeeding regimens are thought to be linked to cycles of stem cell quiescence and activation, using nutrient limitation to remodel metabolism and reset stem cell pools towards a more youthful profile.^10,48,49^

We previously showed that HLH-30/TFEB is a master regulator of ARD, required for proper morphogenesis and survival upon fasting, and tissue regrowth as well as germline regeneration upon refeeding.^16^ Here, we discovered that TFEB is also a key nutrient dependent regulator of germ stem cell dynamics that works via TGFβ signaling, switching stem cells from quiescence to activation likely through a cell non-autonomous mechanism and thereby preventing entry into a senescence-like state. In line with evolutionary conservation, we observed that adipose, liver and muscle tissue of killifish exposed to prolonged fasting show similar upregulation of *tfeb* and *smad4*, which are downregulated upon refeeding, a dynamic regulation that is lost with aging. Furthermore, we extended the key role of TFEB to different diapause settings in mammals, both physiological (embryonic diapause) and pathological (cancer). Interestingly, the TGFβ pathway showed a strong upregulation in the human diapause-like cancer context, in contrast with the model proposed in *C. elegans*. Nevertheless, it has already been described that the TGFβ pathway is wired differently in worms and mammals: a high level of TGFβ promotes worm proliferation/growth^50^, but inhibits proliferation in mammalian cells (though this behavior too depends on cell and signaling context).^51^ Despite seemingly opposite regulation, the two models are reconciled by common output of cytostatic control, supporting the idea that the TFEB-TGFβ axis is essential for diapause regulation. Other studies also provide separate evidence for TFEB or TGFβ signaling in stem cell quiescence, activation, and senescence.^51–54^ Importantly, our studies bring TFEB-TGFβ together in a unified axis regulating these processes.

Besides TGFβ signaling, our screens also identified components of insulin/IGF signaling as mediators of ARD survivorship and recovery. In worms, Insulin-like peptides, similar as TGFβ, are produced mainly in ciliated sensory neurons, and regulate various processes throughout the body via an endocrine mechanism.^55,56^ Accordingly, *hlh-30* loss causes dysregulation of numerous Insulin-like peptides under ARD and refeeding (Figure S5F). We also identified a mutation in *tax-4*/Cyclic Nucleotide Gated cation channel, which also acts within such ciliated sensory neurons to regulate TGFβ^20^ and Insulin-like peptide production.^55^ Interestingly, reduced TGFβ, IIS, and *tax-4* converge on *daf-16*/FOXO to various extents, and *hlh-30* and *daf-16* have overlapping target genes.^57^ Hence, the identification of these additional molecules, is in line with a coherent nutrient-neuron-to-soma relay controlling stem cell dynamics and longevity.

Our model (Figure 6G) provides a powerful conceptual framework to better understand systemic regulation of quiescence/activation, longevity/regeneration, diapause entry/exit beyond *C. elegans*. First, our studies provide seminal evidence linking nutrient regulated TFEB with canonical TGFβ to converge on Notch signaling in a unified regulatory cascade controlling quiescence and activation. Our study concurs with studies linking fasting, TGFβ and insulin/IGF signaling to the regulation of Notch signaling within the GSC and niche.^30,50^ Interestingly, under *AL* conditions, inhibition of *NOTCH* itself (*glp-1*) or laser ablation of germ cells are sufficient to trigger longevity^58^ suggesting that germlineless *glp-1* longevity maps to some extent onto ARD germ cell quiescence. Moreover, Notch signaling is well known to regulate stem cell dynamics in diverse tissues^59^ but has also been linked to mammalian senescence.^60,61^

Second, our studies reveal the basic architecture of how fasting/refeeding can regenerate stem cell pools through a systemic mechanism. As loss of neuronally expressed *daf-7*/TGFβ can bypass *hlh-30*/TFEB collapse, modulating such neural signals might be sufficient to overcome somatic and germline dysfunction. Quite possibly germ cell quiescence/activation is additionally regulated directly by local signaling or nutrient supply. Future studies should pinpoint cellular focus and timing requirements of HLH-30/TFEB action, and unravel nutrient regulation of stem cell quiescence/activation. Notably, systemic regulation in the worm may be conceptually similar to heterochronic parabiosis experiments in mice, where circulating factors from young animals rejuvenate stem cells of older ones.^62^ Systemic factors that regulate mammalian stem cell dynamics and stress response include GDF11, GDF15, and myostatin, which are close relatives of TGFβ.^63,64^ We imagine that some of these secreted factors could also be nutrient regulated via vertebrate TFEB.

Third, our studies may shed unique light on a form of *in vivo* irreversible quiescence resembling cellular senescence. *hlh-30*/*TFEB* mutants in ARD display senescent-like features including failure of stem cell proliferation, enlarged (germ) cell nucleoli, elevated TGFβ and Notch signaling, dysregulated innate immune signaling, mitochondrial dysfunction and DNA damage (Table S1). Furthermore, transcriptomic analysis of biological age indicates *hlh-30* mutants in ARD harbor a premature aging signature, as confirmed physiologically by early onset of a decline in body movement, pharyngeal pumping and SA-β-gal staining. In line with our results, TFEB overexpression in mouse brain mitigates senescence marker and memory deficits in mice^65^, suggesting a potential conserved function. Moreover, TFEB has been recently proposed to play a role in cellular senescence *in vitro*.^66^ We speculate that *hlh-30* mutants cause a disconnect between nutrient availability and growth signaling. Consequently, animals neither fully commit to quiescence nor activation, which could trigger metabolic dysfunction, cellular damage and provoke a senescent-like phenotype. Importantly, however, this senescent-like phenotype can be reversed by modulating TGFβ, a known contributor to mammalian Senescence-Associated Secretory Phenotype (SASP).^67,68^ Conceivably a mismatch between nutrient cues and growth control could also trigger metabolic dysfunction and senescence during vertebrate aging^40^. Indeed, killifish seem to lose their ability to dynamically switch from fasting (high TFEB & SMAD4) to refeeding (low TFEB & SMAD4) in many tissues (Figure 7B). Further study of TFEB-TGFβ axis and other mechanisms underlying ARD may therefore provide unique molecular insights into the dynamics of senescence and its reversal.

In sum, we propose that the TFEB-TGFβ axis has a primordial conserved role in the systemic regulation of diapause states and stem cell quiescence/activation/senescence. Our studies raise several important questions. Among them, what are the time and tissue requirements of this axis in regulating quiescence and activation? What are the critical downstream mechanisms mediating longevity and rejuvenation? Are there specific nutrients, metabolites and metabolic pathways that regulate this process? To what extent do these pathways inform changes in stem cell dynamics during normative aging? Can ARD and recovery bring about a reversal of age-related damage and stimulate somatic and germline rejuvenation? What role does epigenetic remodeling play in these processes? By using the power of a genetically tractable whole animal model, we hope to gain further molecular insight into these processes *in vivo*.

## Acknowledgements

We thank the Caenorhabditis Genetics Center (U. of Minnesota), wormbase.org, the Japanese National Biosource Project for providing strains, Smolikove lab (U. of Iowa) for sharing anti-RAD-51 antibodies, and Antebi lab members for helpful discussion. We thank all students that contributed during internships or rotations (Dylan Aidlen, Logan Rance, Meret Taglinger, Meagan Duncan), Eugene Ballhysa for assistance with gamma irradiation, the bioinformatics core, and the imaging facility for assistance (MPI-AGE). Work in the laboratory of A.A. was supported by the Max Planck Society.

V.R. is recipient of an European Innovative Training Networks H2020-MSCA-ITN-2018 (Healthage - 812830) Marie Curie PhD Fellowship. Work in the laboratory of M.S. was funded by the IRB and “laCaixa” Foundation, and by grants from the Spanish Ministry of Science co-funded by the European Regional Development Fund (ERDF) (SAF2017-82613-R), European Research Council (ERC-2014-AdG/669622), and Secretaria d’Universitats i Recerca del Departament d’Empresa i Coneixement of Catalonia (Grup de Recerca consolidat 2017 SGR 282).

## Author Contributions

A.A., B.G. (worms) and M.S.,V.R. (cells) conceived and designed the study. Investigation, J.M. (EMS mutagenesis screen, genomic & RNAseq, lifespans, *etc*.), T.J.N. (all microscopy data, recovery, *etc*.), B.G. (initial screen, lifespans, recovery, *etc*.), K.L. (*daf-12* data), C.L. (technical assistance), K.K. (Bit age analysis), J.K. (snp mapping), R.R. (killifish data); V.R. (crispr cell screening), Writing – Review & Editing, A.A., B.G., T.J.N., M.S.

## Supplemental Tables

**Table S1:** Hallmarks of cellular senescence.

**Table S2:** Supplementary data will be available upon publication

**Table S3:** Overview of gene ontology biological processes (GOTERM_BP_DIRECT) derived from enrichment analyses of up- or downregulated genes of *hlh-30,* N2 wild-type and *hlh-30 daf-1* under 48 hours of ARD and 12 hours of refeeding.

## Supplemental Figures

**Figure S1: TGF**β **and insulin/IGF signaling in *hlh-30* ARD survival and recovery**

A. and B. Insulin signaling mutants *daf-2(e1368)* (A) and *pdk-1(sa680)* (B) enhance ARD survival of *hlh-30(tm1978)*.

C. Percent reproductive *hlh-30(tm1978);tax-4(p678)* worms that recover after 10 days in ARD upon refeeding. Each dot represents one experiment. Mean ± SEM. One-way ANOVA, ∗∗∗p < 0.001; ∗∗p < 0.01; ns not significant.

D. cGMP signaling mutant *tax-4(p678)* enhances ARD survival of *hlh-30(tm1978).*

Survival curves depict one experiment. Lifespan data and statistics (log-rank tests) are presented in Table S2.

**Figure S2. TGFβ suppression of hlh-30 ARD survivorship and recovery is dependent on downstream transcription factors**

A. *daf-12(rh61rh411)* significantly enhances ARD survival of *hlh-30(tm1978) daf-1(m40).*

B. Percent reproductive worms recovered after 10 days in ARD upon refeeding. Genotypes N2, *hlh-30(tm1978), hlh-30 daf-1(m40),* and *hlh-30 daf-1;daf-12(rh61rh411).* Each dot represents one experiment. Mean ± SEM. One-way ANOVA, ns, not significant.

**Figure S3. HLH-30 regulates TGFβ signaling**

A. *daf-7p::gfp* expression in neurons anterior to the nerve ring in N2 and *hlh-30(tm1978)* background. ASI and another neuron, possibly OLQ (or IL1) at 96 hours of ARD. DIC and fluorescent images of head region of N2. Arrow pointing to neuron. Scale bar 10 μm.

**Figure S4. Regulation of Notch signaling and germ cell phenotypes by *hlh-30* and *daf-1***

A. Representative photomicrographs of distal gonad arms of *sygl-1::3xFlag* in N2 and *hlh-30(tm1978)* worms at 24 hours refeeding. DAPI and ANTI-FLAG antibody staining. Scale bar 10 μm.

B. Representative photomicrographs of germ cell nucleoli of N2, *hlh-30(tm1978)* and *hlh-30 daf-1(m40)* at 96 h of ARD.

**Figure S5. *hlh-30* ARD transcriptome is reversed by *daf-1***

A. Gene ontology enrichment analysis of 8093 DEGs (padj < 0.05) in *hlh-30*/N2 ARD. Top 10 biological processes (DAVID GO BP DIRECT database). Complete GO term list is shown in Table S2.

B. Gene ontology enrichment analysis of 9295 DEGs (padj < 0.05) in *hlh-30*/N2 refeeding. Top 10 biological processes (DAVID GO BP DIRECT database). Complete GO term list is shown in Table S2.

C. Correlation plot of protein coding genes of *hlh-30*/N2 and *hlh-30 daf-1*/*hlh-30* at 12 hours of refeeding. DEGs (padj < 0.05) highlighted in red (genes down in *hlh-30* and reversed by *daf-1,* 1499 DEGs, quadrant 2) or blue (genes up in *hlh-30* and reversed by *daf-1*, 2173 DEGs, quadrant 4). Other DEGs (14022), gray. Simple linear regression line in gray. Equilibration Y = -0.35X - 0.06; R^2^ = 0,27.

D. Gene ontology enrichment analysis of 1499 DEGs down in *hlh-30/*N2 and up in *hlh-30 daf-1/hlh-30* (padj<0.05, quadrant 2) at 12 hours of refeeding. Top 10 biological processes of the DAVID GO BP DIRECT database.

E. Gene ontology enrichment analysis of 2173 DEGs up in *hlh-30/*N2 and down in *hlh-30 daf-1/hlh-30* (padj<0.05, quadrant 4) at 12 hours of refeeding. Top 10 biological processes of the DAVID GO BP DIRECT database.

F. Heat map of *log2(FC)* values of differentially expressed insulin-like peptides in *hlh-30*/N2 at 48 hours of ARD and upon 12 hours of refeeding. padj< 0.05.

**Figure S6. *hlh-30* mutants exhibit senescent-like phenotypes**

A. Representative photomicrographs of dissected distal gonad arms of N2 worms expressing *sygl-1::3xFlag* 1 h post gamma irradiation (60Gy). DAPI and ANTI-FLAG and ANTI-RAD51 antibody staining. Scale bar 10 μm.

B. Representative photomicrographs of SA-β-gal staining in the head region. Genotypes N2, *hlh-30(tm1978),* and *hlh-30 daf-1(m40).*

C. and D. Brood size of self-fertilizing worms refed after 10 days of ARD with bacterial food. Genotypes N2 and *atg-7(bp411)* (C) and N2 and *lgg-2(tm5755)*

(D). Each circle represents the total progeny per worm (total of 100 worms). Pooled data of 2 independent biological replicates. Mean ± SEM from one representative experiment. Mann-Whitney test, ns, not significant.

E. *AL* brood size of self-fertilizing worms. Genotypes N2 and *lgg-2(tm5755)*. Mann-Whitney test

## Methods

### Lead Contact and Materials Availability

Further information and requests for resources and reagents should be directed to and will be fulfilled by the Lead Contact, Adam Antebi. This study did not generate unique reagents or new plasmids. Strains generated in this study are available upon request.

### Experimental Model and Subject Details

Nematodes were cultured using standard techniques at 20 °C on nematode growth medium agar plates with the *Escherichia coli* strain OP50, unless otherwise noted. All strains used in this study are listed in the key resources table. Mutant strains were obtained from the Caenorhabditis Genetics Center (CGC) or National BioResource Project (NBRP) and outcrossed to our N2 wild-type (*C. elegans* variant Bristol). CRISPR–Cas9 mutants were generated by Sunybiotech (https://www.sunybiotech.com). ARD conditions are described below.

## Detailed Methods

### ARD Induction

Eggs, enriched by standard hypochlorite treatment, were grown to the mid-L3 larval stage, assessed by DIC microscopy for migration of the gonad arms. Larvae were collected in M9 buffer and washed four times with M9 buffer. Animals were left to settle for 20 minutes after every wash to allow for expulsion of bacteria from the gut. Afterwards, ca. 600 worms were pipetted onto 3 cm plates containing 4 ml Nematode Growth Medium (NGM) with UltraPureTM agarose (Thermo Fisher Scientific) and 50 μg/ml ampicillin. One day after ARD induction plates were wrapped with parafilm. Worms were maintained for their whole life on one plate at 20 °C, unless noted otherwise.

### Recovery from ARD

ARD worms were transferred to NGM plate seeded with *E. coli* OP50. Worms were monitored every day. Successful exit and recovery from ARD were determined by visual improvements (body size, motility, intestinal coloration, germline growth) and the ability to produce progeny, indicating regeneration of the germline. Worms were categorized as **reproductive adults**, sterile adults or unrecovered ARD worms. For total **brood size** measurement, individual ARD worms were picked to OP50-seeded 3 cm NGM plates, and the progeny number per worm was counted. While producing progeny, the worms were transferred regularly until reproduction had ceased.

### Lifespan Experiments

ARD lifespans were determined by scoring a population of about 600 ARD worms every third day. In one experiment several plates per genotype were scored. Plates that displayed mold or bacteria contaminations were discarded. Day 0 corresponds to the L3 stage (time point of ARD induction).^16^ Lifespan experiments under AL-conditions were performed as described.^69^ Day 0 corresponds to L4 stage.

### *hlh-30* Suppressor Screen

*hlh-30(tm1978)* L4 larvae were mutagenized with 0.5% EMS in M9 buffer according to standard protocols.^70^ Each 10 mutagenized P0 adults were allowed to lay eggs overnight on a 10 cm plate. F1 adults were hypochlorite treated to enrich eggs, and ARD was induced on mid L3 worms of the F2 generation (see above). After 20 days in ARD, *hlh-30* suppressors were transferred to OP50 seeded plates and recovery was monitored (see recovery from ARD). Reproductive worms were singled, resulting in the following strains after re testing: EMS-1 and EMS-7 (test mutagenesis), A1, I4, 37, 39, 40, 43, 51, 67, 70, 71, 87, 104 and 109 (mutagenesis 2).

### Whole Genome Sequencing and Galaxy MiModD Analysis

For genomic sequencing, we prepared genomic DNA from strains listed above with QIAGEN Gentra PureGene tissue kit. Sequence libraries were created using the TruSeq DNA sample prep (Illumina, San Diego, CA). Libraries were sequenced on a HiSeq 2500 (Illumina, San Diego, CA) to generate 150 bp paired end reads. Library preparation and sequencing was performed by the Max Planck Genome Center (Cologne, Germany, https://mpgc.mpipz.mpg.de/home/). Sequencing data was analyzed using Galaxy software. The WS220/ce10 *C. elegans* assembly was used as reference genome for annotation.

SNP-based mapping was performed on mutant strains A1, 39, 40, 43, 71, 109 by crossing them to *hlh-30(tm1978)* in the Hawaiian strain CB4856 background. F1 generation was bleached and ARD was induced on the F2 generation. After 20 days, ARD worms were transferred to OP50 seeded plates, recovered worms were singled on 6 cm plates and progeny was pooled for genomic DNA preparation. Pooled DNA was sequenced on an Illumina HiSeq platform (paired-end 150[nt). Mutations were identified with MiModD software (https://celegans.biologie.uni-freiburg.de/?page_id=917). WS220/ce10 *C. elegans* assembly was used as reference genome for annotation.

Causative mutations were confirmed by testing existing deletions or CRISPR– Cas9 designed bp change of identified genes in the *hlh-30* background on ARD recovery.

### Dauer Assay

To determine dauer formation, worms were age synchronized by allowing 20 worms to lay eggs for 4 hours. 100 eggs were picked to a fresh plate and incubated at 25 °C for 48 hours. To assess dauer formation, dauer characteristics such as constricted pharynx and long and thin body shape were used. To study whether *daf-3(syb2718)* enter dauer in a dominant/recessive fashion we took advantage of the recessive dumpy phenotype as a marker for heterozygosity. For this experiment *daf-3(syb2718);dpy-1* and control worms were crossed with wild-type males. F1 cross progeny eggs were shifted to 25°C for 48 hours and heterozygous worms (no dumpy phenotype) were scored for dauer entry.

### Worm Size Measurements

Images of worms were taken with Zeiss Axio Imager Z1 or Leica M165 FC microscope. Body length was determined using Image J. At least 25 worms were analyzed per genotype.

### RNA Sequencing *C. elegans*

For RNA sequencing, total RNA was prepared from at least 3000 worms per genotype, using the RNeasy Mini Kit (QIAGEN). Four independent biological replicates were prepared. polyA+ mRNA was isolated using NEBNext Poly(A) mRNA Magnetics Isolation Module (New England Biolabs), RNA-seq libraries were prepared with the NEBNext Ultra Directional RNA Library Prep Kit for Illumina (New England Biolabs), quantified by fluorometry, immobilized and processed onto a flow cell with a cBot (Illumina) followed by sequencing-by-synthesis with TruSeq v3 chemistry on a HiSeq2500 at the Max Planck Genome Center (Cologne, Germany).

Reads were quality trimmed with Flexbar version 2.5, then mapped to the reference genome (WBcel235.80) using hisat2 version 2.0.4. Respective assemblies were merged with cuffmerge, version 2.2.1, differential gene expression analysis was performed with Cuffquant version 2.2.1 and Cuffdiff version 2.2.1.

GO annotation and enrichment was performed using DAVID bioinformatics resource database analysis via Flaski web app for data analysis and visualization^71^ developed by the Bioinformatics Core Facility of the Max Planck Institute for Biology of Ageing, Cologne, Germany.

### RNA Sequencing killifish

All experiments were performed on adult (young 6-8 weeks, and old 18-20 weeks old) African turquoise killifish *Nothobranchius furzeri* laboratory strain GRZ-AD. Adult fish were single-housed in 2.8 L tanks from the second week of life. Water parameters were pH 7-8, kH 3-5, and T 27.5 °C., 12 h of light and 12 h of darkness and fed with 10 mg of the dry pellet (BioMar INICIO Plus G) and Premium Artemia Coppens® twice a day. The fish were either fasted for 72 h or fasted for the same amount of time and refed for 24 h and then euthanized. To reduce variability due to circadian rhythms, fish were sacrificed all at once within two hours in the early afternoon. Harvested tissues were snap-frozen in liquid nitrogen and stored at –80 °C. RNA extraction of all samples was done at the same time. 1 μg of total RNA was used for library preparation. The sequencing was performed on the Illumina HiSeq4000 sequencing system (∼50 million reads per sample) using a paired-end two × 100 nt sequencing protocol. After removing rRNA and tRNAs, reads were pseudo-aligned to the reference genome (Nfu_20140520) using Kallisto (0.45.0). Pair-wise differential gene expression was performed using DESeq2 (1.24.0). Animal experimentation was approved by “Landesamt für Natur, Umwelt und Verbraucherschutz Nordrhein-Westfalen”: 81-02.04.2019.A055.

### Biological age prediction of worms

Biological age prediction of worms was carried out using transcriptome-based aging clock, BiT age.^36^ Source code was downloaded from https://github.com/Meyer-DH/AgingClock. Counts-per-million normalized RNA-seq reads for each genotype were used as input for BiT age analysis.

### Imaging and Image Analysis

For live imaging, worms were anaesthetized in 0.1% sodium azide. Image analysis was performed on 2D images using Fiji software.^72^ ***hlh-30::mNeonGreen*** nuclear localization in ASI neurones was determined at 2h and 48h of ARD and after refeeding 48h ARD worms for 1h, 4h, 6h, or 24h using Zeiss Axio Imager Z1 microscope. Nuclear localization ratios were determined by measuring the fluorescence intensity in the nuclear and the cytosolic region of the ASI neurons. ***daf-7p::gfp*** expression was determined at 48h, 72h and 96h of ARD and 48h of ARD refeeding in one ASI neuron per worm using Zeiss Axio Imager Z1 microscope. *daf-7p::gfp* expression in the OLQ was determined after 96h of ARD. ***daf-1p::gfp*** expression in the DTC was determined at 48h of ARD and 48h of refeeding 48h ARD worms using Zeiss Axio Imager Z1 microscope. Whole body *daf-1p::gfp* expression was monitored with the COPAS Biosorter (Union Biometrica, setting: green 450). ***lag-2p::mCherry-PH*** expression was determined in 48h old ARD worms. Worms were imaged with the Leica SP8 confocal microscope. Z-stacks of whole DTCs (z-stack size 0.3µm) were captured. The sum-projection of the DTC z-stacks was used to measure the total fluorescence intensity of the DTC cap structure. Maximal DTC length was determined by measuring the length of the longest DTC extension from distal to proximal using the *lag-2p::mCherry-PH* construct to color the DTC. Germ cell nucleolar area was determined from photos of the gonad taken with a Zeiss Axio Imager Z1 microscope (DIC contrast). Per genotype 15 gonad arms from different animals were analyzed, scoring the area of each 3-5 nuclei located in the vicinity of the DTC. For different **Immunofluorescence** analysis, germline dissections and staining were performed as described previously.^73^ Briefly, ARD and refed ARD worms were anaesthetized in 200mM levamisole, dissected, fixed in 2% formaldehyde and post-fixated in 100% Methanol at -20 for 10 min. Fixates were blocked in 30% Goat serum (Cell Signaling #5425S) or 1% BSA and stained with different primary antibody (Phospho-Histone H3, 1:150, Cell Signaling #9706S; ANTI-FLAG, 1:1000, Sigma-Aldrich #F3165, ANTI-RAD51, 1:1000 a gift from Smolikove lab) over night. Secondary antibody staining (Alexa Fluor^TM^ 488 #A-11008, Alexa Fluor^TM^ 594 #A-11005, 1:400) was followed by fixate embedding in Fluoromount-G^TM^ (Invitrogen #00-4959-52) containing DAPI. Z-stacks of whole gonad arms (z-stack size 0.3µm) were captured using the Leica SP8 or Leica DLS confocal microscope. Images analysis was performed on z-projections in Fiji^72^ and for the cell cycle analysis using 3D images viewer of Imaris.

### Senescence-Associated **β**-Galactosidase assay

This assay was performed with equal amounts of 48h old ARD worms after freeze-cracking using a Senescence β-Galactosidase Staining Kit (Cell Signaling, #9860) according to manufactures instructions. Image analysis was performed in Fiji^72^ and color values were generated on RGB images.

### Cell culture procedure

Embryonic Stem cells (ES) cells were grown on gelatin 1% with ES medium (EM): Dulbecco’s modified Eagle’s medium (DMEM, high glucose, Life Technologies) supplemented with 15% foetal bovine serum (FBS, Gibco), penicillin/streptomycin (100 U/mL), LIF (1000 U/ml), 0.1 mM non-essential amino acids, 1% glutamax, and 55 mM β-mercaptoethanol). Diapause was induced by 200nM INK128 (S2811, SelleckChem).

SK-Mel-103 cells were grown in Dulbecco’s modified Eagle’s medium (DMEM, high glucose, Life Technologies) supplemented with 10% foetal bovine serum (FBS) (Gibco) and penicillin/streptomycin (100 U/mL). The diapause-like state was induced by 100nM INK128.

### Genome-wide sgRNA screening

To perform the genome-wide screen, we used a CRISPR/Cas9 system previously described.^43^ Embryonic stem cells were generated from mice ubiquitously expressing Cas9 under control of the endogenous Cola1a locus and a tetracycline responsive operator transgene; reverse tetracycline-controlled transactivator synthesis is under control of the endogenous ROSA26 locus. ESCs from these mice were infected with a lentiviral library encoding short guide RNAs (sgRNAs) targeting 19,150 mouse genes, with approximately five independent sgRNAs per gene. Addition of doxycycline to the ESCs causes inducible expression of the Cas9 enzyme, which induces editing of the sgRNA targets.

Diapause was induced in ESCs through the mTOR/PI3K inhibitor INK128 at 200nM. After 5 days, half of the proliferating cells and half of the diapaused cells were sequenced to know the initial diversity of the sgRNAs library. Doxycycline was added to the remaining cells for 3 additional days to induce sgRNA editing before sequencing.

Genomic DNA was isolated from cell pellets using a genomic DNA isolation kit (Blood & Cell Culture Midi kit (Qiagen). After gDNA isolation, sgRNAs were amplified and barcoded by PCR as in Shalem et al., to amplify the DNA fragment containing sgRNA sequences. PCR products were sequenced on a HiSeq 4000 instrument (Illumina) at 50bp reads to a depth of 30M reads per sample.

Reads were preprocessed by removing adapters using Cutadapt v4.1^74^ with parameters “-e 0.2 -a GTTTTAGAGCTAGAAATAGCAAGTTAAAATA -m 18”. Next sequences were trimmed to length 19 to match that of the probes. An artificial genome was created for alignment with Bowtie v0.12.9^75^ using the probes’ sequences and the function bowtie-build with default parameters. Finally, reads were aligned to this genome with parameters -S -t -p 20 -n 1 -l 19.

Read counts were imported into R^76^ by reading the sam files and counting the number of occurrences of each probe. The resulting count matrix was normalized using the rlog function from the DESeq2 v1.34.0^77^ R package.

A linear model with random and fixed effects was fitted to the normalize probe data for each gene. The condition was used as a fixed covariable while the guides were included as random effects whenever there was more than one. The model was fit with the function lmer from the lme4^77^ package or with the native R lm function if there was only one probe for a given gene. Contrast coefficients and p-values were computed using the glht function from the multcomp^78^ package without any p-value adjustment.

GO gene set collections were downloaded from the Gene Ontology knowledgebase^79^. Genes quantified in the microarray study were annotated according to the Broad Hallmark.^80^

Functional enrichment analyses were performed using a modification of ROAST^81^, a rotation-based approach implemented in the Limma R package^82^, which is especially suitable for small experiments. Such modifications were implemented to accommodate the proposed statistical restandardization^83^ in the ROAST algorithm, which enables its use for competitive testing.^84^ The MaxMean^83^ statistic was used for testing gene-set enrichment of the different gene collections. For each gene, the most variable guide within each gene was used in these analyses (median absolute deviation). The results of these analyses were adjusted by multiple comparisons using the Benjamini-Hochberg False Discovery Rate method.^85^ All these was performed using the functions of the roastgsa^86^ R package.

### siRNA treatment

siRNA for TFEB was purchased from siTOOLs. siRNAs (non targeting and TFEB) have been used at 3nM final concentration. Both proliferating and diapause-like SK-Mel-103 have been transfected with siRNAs for 5 days before performing viability assays. Lipofectamine reagent RNAiMAX (Cat. 13778075) was used at 2uL per mL to perform the transfection.

Cell viability has been measured using Cell Titer Glo Luminescent cell viability assay (Promega). Raw data were acquired by measuring luminescence in a VICTOR Multilabel Plate Reader (Pelkin Elmer).

### Quantification and Statistical Analysis

The statistical tests performed in this study are indicated in figure legends and method details. Data are represented as mean ± SEM as stated in the figure legends. Number of replicates and animals for each experiment are enclosed in their respective figure legends and or method details.

### Data and Code Availability

RNA-seq data will available upon publication.

## Resources Table

| REAGENT or RESOURCE | SOURCE | IDENTIFIER |
| --- | --- | --- |
| Antibodies |  |  |
| Phospho-Histone H3 (Ser10) Mouse mAb | Cell Signaling | #9706 |
| Fluoromount-G<sup>TM</sup> | Invitrogen | #00-4959-52 |
| Goat serum | Cell Signaling | #5425S |
| ANTI-FLAG | Sigma-Aldrich | #F3165 |
| Alexa Fluor<sup>TM</sup> 488, | Invitrogen | #A-11008 |
| Alexa Fluor<sup>TM</sup> | Invitrogen | 594 #A-11005 |
| Bacterial Strains |  |  |
| *E. coli*: Strain OP-50 | Caenorhabditis Genetics Center | WormBase: OP50 |
| **Biological Samples** |  |  |
| SK-Mel-103 | ATCC |  |
| KH2 embryonic stem cells | Oscar Fernandez-Capetillo's laboratory |  |
| **Chemicals, Peptides, and Recombinant Proteins** |  |  |
| UltraPure™ agarose | Thermo Fisher Scientific | Cat#16500-500 |
| C<sub>1</sub>-BODIPY 500/510 C<sub>12</sub> | Invitrogen™ | Cat#D3823 |
| INK128 (Sapanisertib) | SelleckChem | Cat#S2811 |
| **Critical Commercial Assays** |  |  |
| RNAeasy Mini Kit | QIAGEN | Cat#74104 |
| NEBNext Poly(A) mRNA Magnetis Isolation Module | New England Biolabs | Cat#E7490S |
| NEBNext Ultra Directional RNA Library Prep Kit for Illumina | New England | Cat#E7420S |
| Senescence β-Galactosidase Staining Kit | Cell Signaling | Cat#9860 |
| CellTiter-Glo Luminescent Cell Viability Assay | Promega | Cat#G7571 |
| **Deposited Data** |  |  |
| RNA seq data | AA lab | NCBI GEO accession number: XXX |
| **Experimental Models: Organisms/Strains** |  |  |
| *C. elegans*: Strain N2 wild isolate | CGC | WormBase: N2 |
| *C. elegans*: *hlh-30(tm1978)* IV. | AA lab | Strain AA3658 |
| *C. elegans*: *daf-1(m40)* IV. | CGC | Strain DR40 |
| *C. elegans*: *daf-7(e1372)* III. | CGC | Strain CB1372 |
| *C. elegans*: *pdk-1(sa680)* X. | CGC | Strain JT9609 |
| *C. elegans*: *tax-4(p678)* III. | CGC | Strain PR678 |
| *C. elegans*: *daf-2(e1368)* III. | AA lab | Strain AA4401 |
| *C. elegans*: *daf-2(e1370)* III. | AA lab | Strain AA5311 |
| *C. elegans*: *daf-3(syb2718)* X. | SunyBiotech | Strain PHX2718 |
| *C. elegans*: *daf-3(syb2718)* X; *dpy-1(e1)* III. | AA lab | Strain AA5417 |
| *C. elegans*: *daf-3(e1376)* X. | CGC | Strain CB1376 |
| *C. elegans*: *daf-5(e1386)* II. | AA lab | Strain AA4397 |
| *C. elegans*: *daf-16(mgDf50)* I. | AA lab | Strain AA60 |
| *C. elegans*: *daf-1(m40) hlh-30(tm178)* IV. | AA lab | Strain AA5029 |
| *C. elegans*: *daf-7(e1372)* III; *hlh-30(tm178)* IV. | AA lab | Strain AA5030 |
| *C. elegans*: *pdk-1(sa680)* X; *hlh-30(tm1978)* IV | AA lab | Strain AA5033 |
| *C. elegans*: *tax-4(p678)* III; *hlh-30(tm1978)* IV. | AA lab | Strain AA5325 |
| *C. elegans*: *daf-2(e1370)* III; *hlh-30(tm1978)* IV. | AA lab | Strain AA5450 |
| *C. elegans*: *daf-3(e1376)* III; *hlh-30(tm1978)* IV. | AA lab | Strain AA5339 |
| *C. elegans*: *daf-3(syb2718)* X; *hlh-30(tm1978)* IV. | AA lab | Strain AA5252 |
| *C. elegans*: *daf-12(rh61rh411)*X; *hlh-30(tm1978)* IV | AA lab | Strain AA5289 |
| *C. elegans*: *daf-1(m40) hlh-30(tm178)* IV; *daf-3(e1376)* X. | AA lab | Strain AA5034 |
| *C. elegans*: *daf-1(m40) hlh-30(tm178)* IV; *daf-5(e1386)* II. | AA lab | Strain AA5036 |
| *C. elegans*: *daf-1(m40) hlh-30(tm178)* IV; *daf-16(mgDf50)* I. | AA lab | Strain AA5035 |
| *C. elegans*: *daf-1(m40) hlh-30(tm178) IV*; *daf-12(rh61rh411)X* | AA lab | Strain AA5466 |
| *C. elegans*: *daf-7(e1372) III*; *hlh-30(tm178) IV*; *daf-3(e1376) X*. | AA lab | Strain AA5037 |
| *C. elegans*: *daf-7(e1372) III*; *hlh-30(tm178) IV*; *daf-5(e1386) II*. | AA lab | Strain AA5038 |
| *C. elegans*: *mEx40[rol-6(su1006) daf-1p::GFP]*. | Riddle lab | Strain DR1767<br>*daf-1p::gfp* |
| *C. elegans*: *hlh-30(tm1978)*; *mEx40[rol-6(su1006) daf-1p::GFP]*. | AA lab | Strain AA5553 |
| *C. elegans*: *mls7(daf-7p::GFP::daf-7 3«UTR)*. | Riddle lab | Strain DR2021<br>*daf-7p::gfp* |
| *C. elegans*: *hlh-30(tm1978) IV*; *mls7(daf-7p::GFP::daf-7 3«UTR)*. | AA lab | Strain AA4369 |
| *C. elegans*: N2; *nals37[pGC457 Plag-2(3kb)::mCherryPH::let-858(3')*; *unc-119(+)*]. | Hubbard lab | Strain GC1038<br>*lag-2p::mCherry-PH* |
| *C. elegans*: *hlh-30(tm1978) IV*; *nals37[pGC457 Plag-2(3kb)::mCherryPH::let-858(3')*; *unc-119(+)*]. | AA lab | Strain AA5336 |
| *C. elegans*: *daf-1(m40) hlh-30(tm1978) IV*; *nals37[pGC457 Plag-2(3kb)::mCherryPH::let-858(3')*; *unc-119(+)*]. | AA lab | Strain AA5338 |
| *C. elegans*: *hlh-30(syb809[hlh30::mNeonGreen]) IV*. | SunyBiotech | Strain PHX809<br>*hlh-30::mNeonGreen* |
| *C. elegans*: *daf-3(ot875[daf-3::GFP::3xFlag]) X*. | CGC | Strain OH14892<br>*daf-3::gfp* |
| *C. elegans*: CB1386. | CGC | Strain CB4856<br>Hawaiian strain |
| *C. elegans*: *hlh-30(tm1978)* in CB1386. 8 back crosses. | AA lab | Strain AA4398 |
| *C. elegans*: *sygl-1(am307[3xFLAG::sygl-1]) I*. | CGC | Strain WU1770 |
| *C. elegans*: *sygl-1(am307[3xFLAG::sygl-1]) I*; *hlh-30(tm1978) IV* | AA lab | Strain AA5415 |
| *C. elegans*: *atg-7(bp411) IV* | Melendez lab | Strain QU190 |
| *C. elegans*: *lgg-2(tm5755) IV* | AA lab | Strain AA4371 |
| Killifish (*Nothobranchius fuzeri*) | AA lab | Strain: GRZ-AD |
| **Software and Algorithms** |  |  |
| Flaski | Bioinformatics Core Facility of the Max Planck Institute for Biology of Ageing | N/A |
| Galaxy platform | https://galaxyproject.org/ | N/A |
| MiModD | http://www.celegans.de/en/mimodd |  |
| LAS X | Leica Microsystems | N/A |
| Zen 2.3 | Zeiss Microscopy | N/A |
| Flow Pilot | Union Biometrica | N/A |
| ImageJ | NIH | N/A |
| Imaris |  |  |
| Photoshop and Illustrator | Adobe | N/A |
| Excel | Microsoft | N/A |
| Prism | GraphPad | N/A |
| R (survival) | (Therneau, 2015) | N/A |
| Flexbar v2.5 | <sup>87</sup> | N/A |
| Hisat v0.1.6-beta | <sup>88</sup> | N/A |
| StringTie v1.04 | <sup>89</sup> | N/A |
| Cufflinks v2.2.1 | <sup>90</sup> | N/A |
| **Other** |  |  |
| Sunybiotech: CRISPR–Cas9 mutants | https://www.sunybiotech.com |  |

WB, worm base. CGC, Caenorhabditis Genetics Center. AA lab, Adam Antebi lab, N/A, not applicable.

