## Supplementary Table 1 for "A primordial TFEB-TGFβ signaling axis systemically regulates diapause and stem cell longevity"

**Table S1: HLH-30 protects worms in ARD & refeeding from cellular senescence features.**

| **Hallmarks of cellular senescence** | ***hlh-30* ARD & refeeding** |
|---|---|
| Non reversible cell-cycle arrest | ✓ |
| Activation of cell-cycle inhibitors p21 (CDKN1a/CIP1) and p16 (CDKN2a/INK4) and others | nd |
| Accumulation of DNA damage (persistent DDR, DNA damage foci) | ✓ |
| Senescence-associated heterochromatic foci (SAHFs) | nd |
| Altered cell morphology | ✓ |
| Increased lysosomal mass & activity / Senescence-associated β-galactosidase (SA-βGal) activity | ✓ |
| Altered metabolism | ✓ |
| Mitochondrial dysfunction | (Gerisch et al., 2020) |
| Increased ROS | nd |
| Reduced LaminB1 in nuclear membrane | nd |
| Apoptosis exclusion | nd |
| Senescence-associated secretory phenotype (SASP) | ✓ |
| Chronic inflammation | ✓ |
| Altered NOTCH1 signaling | ✓ |
| Higher mTORC1activity | |
| Autophagy dysfunction | (Gerisch et al., 2020) |
| Dysregulated unfolded protein response | ✓ |

✓ shown in this publication.

Selected hallmarks of senescence adapted from (Gonzalez-Gualda, Baker, Fruk, & Munoz-Espin, 2021).

Gerisch, B., Tharyan, R. G., Mak, J., Denzel, S. I., Popkes-van Oepen, T., Henn, N., & Antebi, A. (2020). HLH-30/TFEB Is a Master Regulator of Reproductive Quiescence. *Dev Cell, 53*(3), 316-329 e315. doi:10.1016/j.devcel.2020.03.014
Gonzalez-Gualda, E., Baker, A. G., Fruk, L., & Munoz-Espin, D. (2021). A guide to assessing cellular senescence in vitro and in vivo. *FEBS J, 288*(1), 56-80. doi:10.1111/febs.15570
