## Supplementary figures for "A primordial TFEB-TGFβ signaling axis systemically regulates diapause and stem cell longevity"

### Supplement figure 1

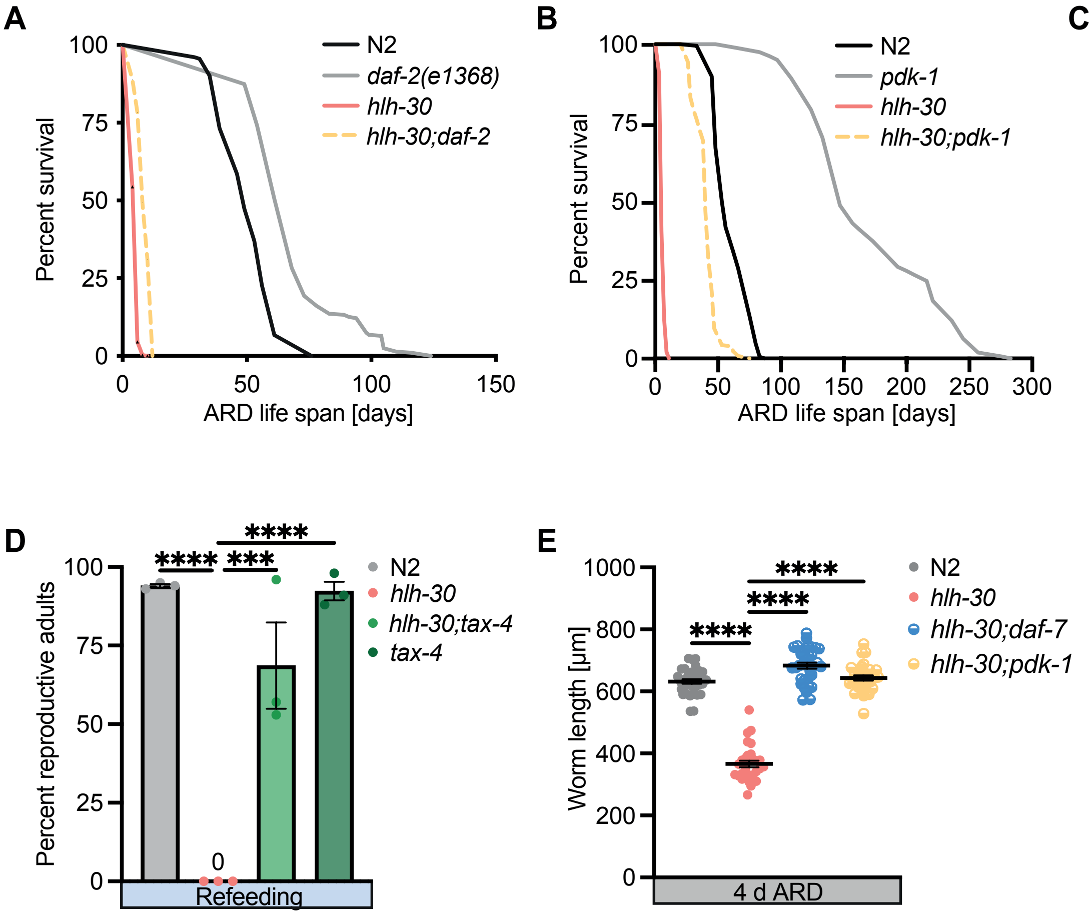

### Supplement figure 2

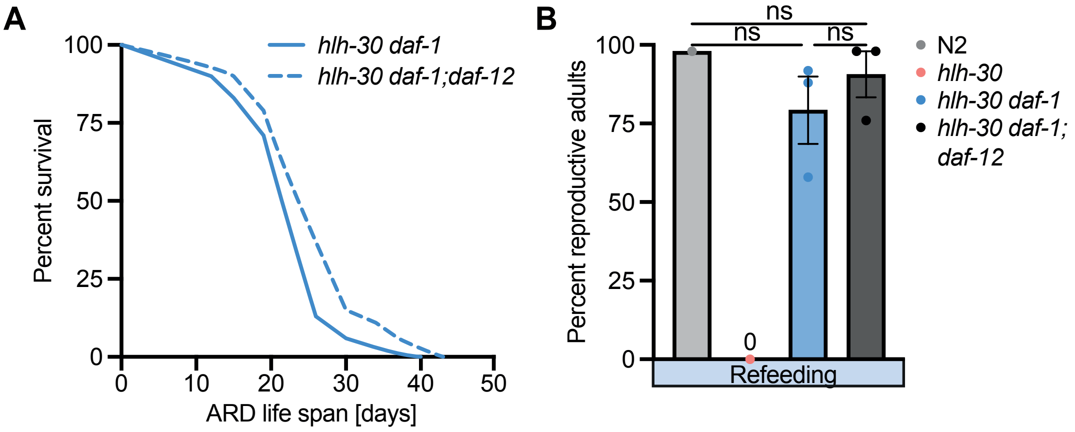

**Supplement figure 3**

**A**

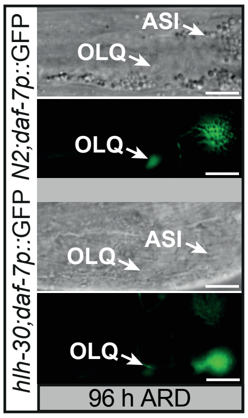

### Supplement figure 4

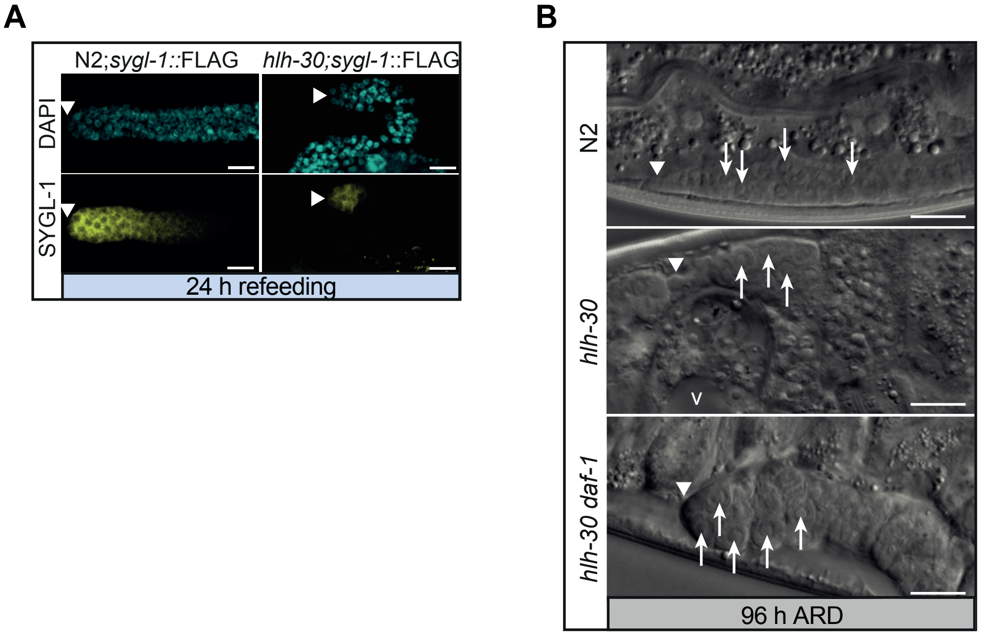

### Supplement figure 5

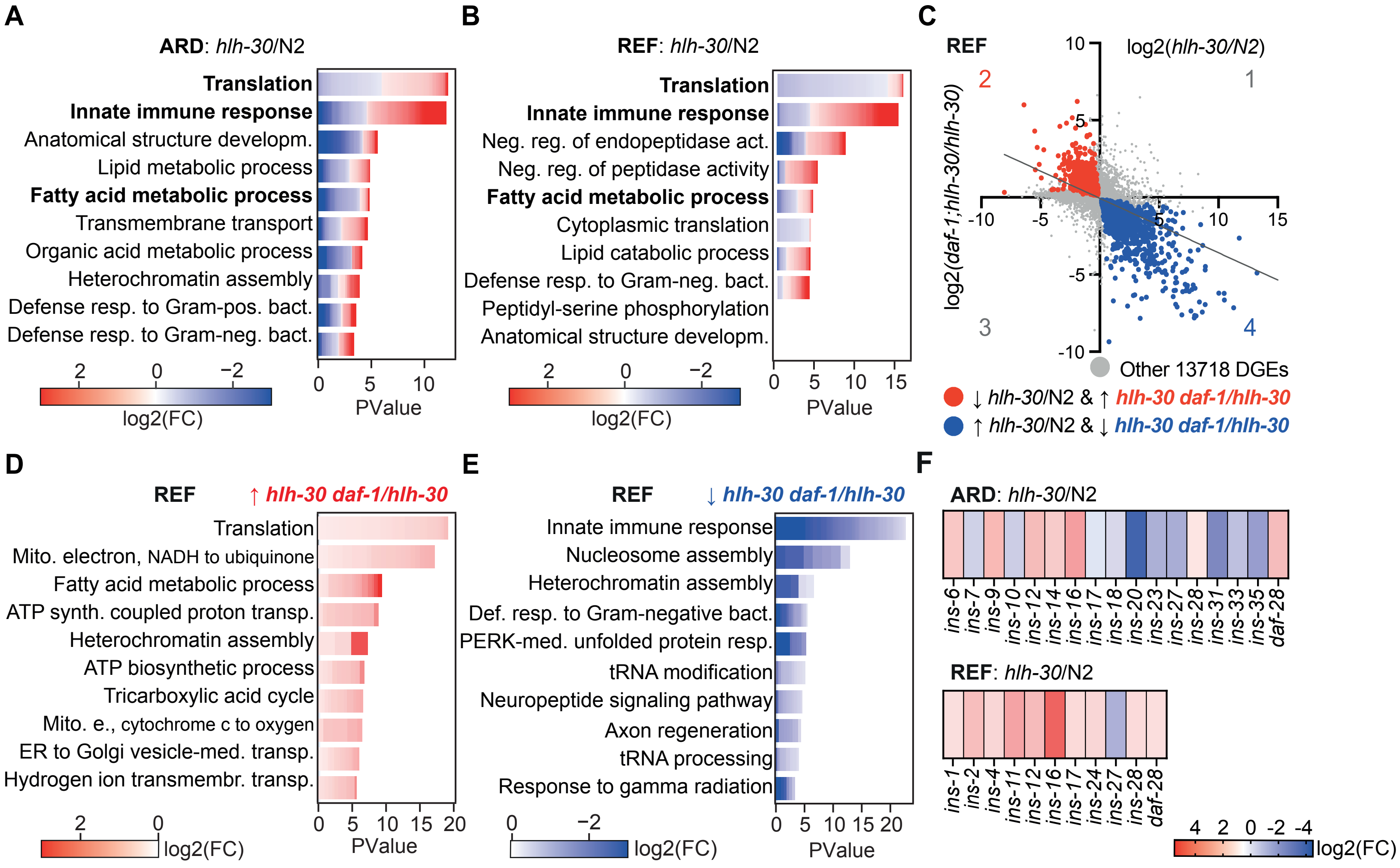

### Supplement figure 6

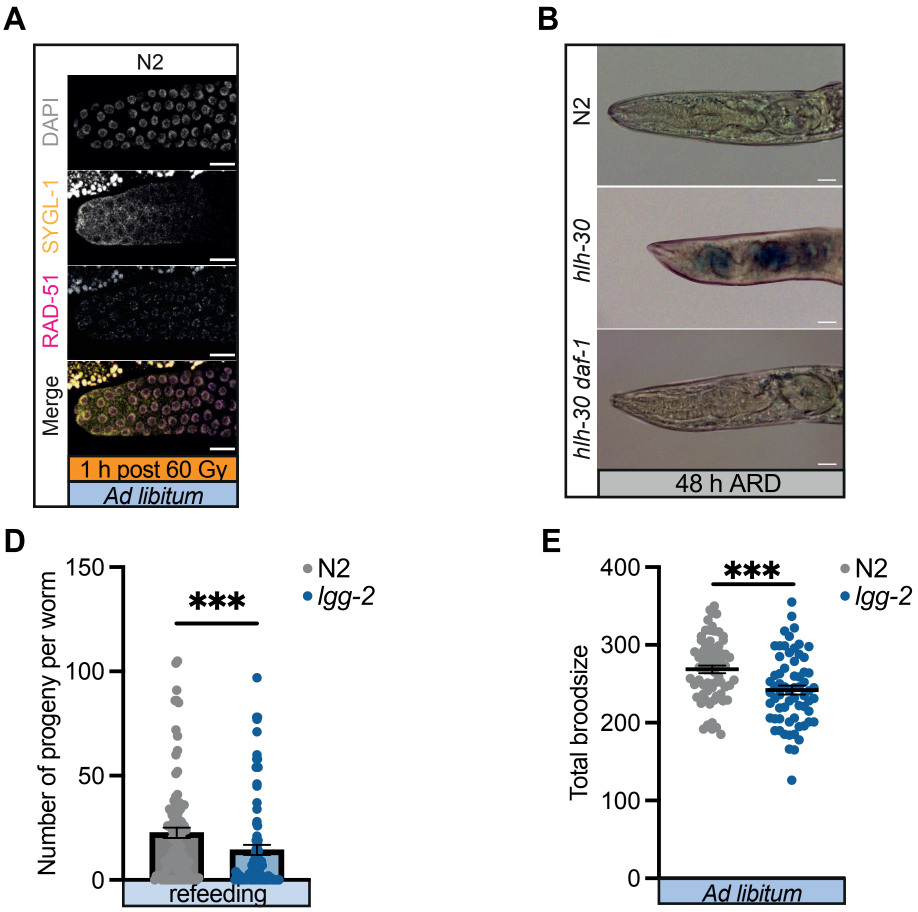
